# Dependencies in heterogeneous, lineage plastic patient–derived prostate cancer organoids revealed through integrated single–cell multiomics and CRISPR screening

**DOI:** 10.64898/2026.05.07.723570

**Authors:** Samir Zaidi, Maren Büttner, Weiran Feng, Caitlin Baxter, D. Henry Kates, Tiago Paiva Prudente, Diana Zakharova, Sanjoy Mehta, Subhiksha Nandakumar, Chen Khuan Wong, Hojin Lee, Pierre–Jacques Hamard, Wenfei Kang, Jungmin Choi, Ronan Chaligné, Robert Cohen, Yu Chen, Ari Firestone, Zhenghao Chen, Charles L. Sawyers

## Abstract

Lineage plasticity and tumor heterogeneity limit the effectiveness of targeted therapies, yet the functional dependencies used to nominate therapeutic targets are often derived from homogeneous systems that fail to capture this complexity. Here, we establish a framework to resolve state–specific genetic vulnerabilities by integrating single–cell multiomics (RNA and ATAC) with pooled CRISPR–Cas9 screening across a large panel of patient–derived organoids (PDOs) from castrate–resistant prostate cancer (CRPC) and neuroendocrine prostate cancer (NEPC). We generate a single–cell multiome atlas spanning >190,000 cells across 22 PDOs, defining seven lineage states—including intermediate and plastic populations not resolved by bulk profiling—and demonstrate that these lineage programs robustly classify independent transcriptomic datasets from prostate cancer patient tumors. By systematically coupling this atlas to subtype–resolved CRISPR screens, we construct a functional dependency map linking cell state in heterogeneous 3D human tumor models. We show that intratumoral heterogeneity fundamentally reshapes the interpretation of gene essentiality, whereby gene–level depletion reflects the composite behavior of co–existing subpopulations, and identify a general principle in which resistant “limiting” populations disproportionately determine aggregate fitness effects. This framework reveals both canonical and previously unrecognized lineage–restricted dependencies within highly plastic tumor and NEPC states, including a therapeutically targetable dependency on the aryl hydrocarbon receptor (AHR) in a novel hybrid stem–like/ASCL1 population. Together, these data establish an extensive multi–dimensional prostate cancer resource, identify novel lineage–resolved biology, and provide a generalizable strategy for interpreting functional genomics in heterogeneous human tumors.

## Introduction

Tumor heterogeneity is a defining feature of advanced malignancies and a major driver of therapeutic resistance^1^. Despite extensive transcriptional and epigenetic profiling, the functional vulnerabilities of distinct tumor cell states—and how these vulnerabilities are shaped by their coexistence within heterogeneous tumors—remain poorly resolved^2–4^. A central challenge is that functional dependencies are typically inferred from bulk or homogeneous systems, implicitly assuming uniform dependency landscapes, despite growing evidence that intratumoral heterogeneity profoundly alters both the strength and interpretability of such signals^5^. This is particularly evident in cancers treated with targeted therapies, where selective pressure promotes adaptive diversification of cell states rather than durable eradication^6^. A key mechanism underlying this process is lineage plasticity, the ability of tumor cells to reversibly alter cellular identity in response to environmental or therapeutic stress, which is associated with poor clinical outcomes across multiple tumor types^7^.

Prostate cancer represents a paradigmatic model of therapy–induced lineage plasticity. Following treatment with androgen receptor pathway inhibitors (ARPIs), tumors frequently transition from androgen–dependent adenocarcinoma toward alternative lineage states, culminating in its most extreme form into neuroendocrine prostate cancer (NEPC), a highly aggressive and treatment–resistant disease^8, 9^. Recent profiling studies have broadly classified prostate cancer heterogeneity into androgen–dependent, WNT–driven, and neuroendocrine lineages, as well as a fourth lineage with AR–low or AR-negative states—often described as stem–like, poorly differentiated, or lineage–ambiguous^10^. However, the latter populations feature intermediate and plastic states, which exhibit substantial internal heterogeneity, often co–exist within tumors, and are poorly resolved by bulk analyses,^2^ leaving their regulatory programs and therapeutic vulnerabilities incompletely understood.

We and others have identified a distinct plastic AR–low state driven by JAK/STAT and FGFR signaling, that can be therapeutically modulated^8, 11^. Epigenetic regulators such as *EZH2* and *NSD2* have been implicated in maintaining AR–low and NEPC states, highlighting a potential path forward for lineage–directed therapies^12, 13,14^. Despite these advances, clinical translation remains limited, in part because current approaches fail to adequately resolve the diversity of plastic states or to match therapies to specific tumor subpopulations.

Several fundamental questions remain: (1) which transcription factors and druggable genes sustain heterogeneous plastic states, (2) can these dependencies be selectively targeted without inducing compensatory transitions, and (3) how does intratumoral heterogeneity shape functional dependency landscapes? Addressing these challenges has been difficult due to limitations of bulk profiling and the inability of most functional genomics platforms to directly link cell state with genetic dependency in the context of heterogeneous model systems. Additionally, homogenous models of AR–low and NEPC states are underrepresented in resources such as DepMap projects highlighting the need for more realistic models that faithfully recapitulate these clinically important lineages.

Patient–derived organoids (PDOs) preserve tumor heterogeneity and co–existing lineage states, making them well–suited to study lineage plasticity. Here, we address the limited availability of functional genomics resource across prostate cancer lineage states by integrating single–cell transcriptomic and chromatin accessibility profiling with pooled CRISPR–Cas9 screening across a large panel of human prostate cancer PDOs. We construct a single cell multiome atlas spanning 22 PDOs and define seven major lineage states, including intermediate and plastic populations obscured in bulk analyses. Leveraging this framework, we perform subtype–resolved CRISPR screens to map functional dependencies across lineage states. By linking gene depletion to transcriptional and chromatin states, we uncover state–specific vulnerabilities and demonstrate how intratumoral heterogeneity alters the interpretation of functional screens. Using this framework, we identify both established and previously unrecognized lineage–restricted dependencies, including actionable vulnerabilities within highly plastic states and a context–dependent dependency on the aryl hydrocarbon receptor (AHR) in prostate cancer. Together, these data establish a comprehensive multi–dimensional resource linking cell state, regulatory programs, and functional dependencies, and provide a generalizable framework for targeting lineage plasticity in advanced prostate cancer and other solid tumors.

## RESULTS

### Single Cell Multiome Analyses of PDO Atlas

To resolve cell heterogeneity and regulatory dynamics beyond the limits of bulk tumor profiling, we performed paired single–cell gene expression and chromatin accessibility (sc–multiome) sequencing on 22 PDOs generated from castrate–resistant prostate cancer (CRPC) and NEPC tumors. Seventeen of these PDOs had been previously classified by bulk RNA– and ATAC–sequencing into four major subtypes: AR (androgen–receptor), SCL (stem–cell–like), WNT (WNT signaling), and NEPC^10^. While this classification scheme has been informative, we sought to determine whether sc–multiome profiling could refine lineage definitions, uncover regulatory states not captured by gene expression alone, and quantify heterogeneity within and across PDOs.

UMAP projection of 195,180 profiled cells revealed substantial heterogeneity across 22 PDOs, which largely segregated by sample (Fig 1A). Annotation by bulk–defined subtypes and cell cycle states were not primary drivers of this separation (Fig S1A and S1B). We therefore applied a multi–stage sub–clustering strategy to identify shared transcriptional and regulatory programs across PDOs. Briefly, we first performed PDO–specific Leiden clustering independently for RNA and ATAC modalities to avoid cross–sample averaging, followed by annotation using established lineage markers (e.g., *KLK3, TP63, NKD1, ASCL1,* and *NEUROD1*; Table S1). Subtypes were retained if their top 20 marker genes or peaks were differentially enriched relative to the remaining subtypes across the atlas (Fig S1C).

**Figure 1.**
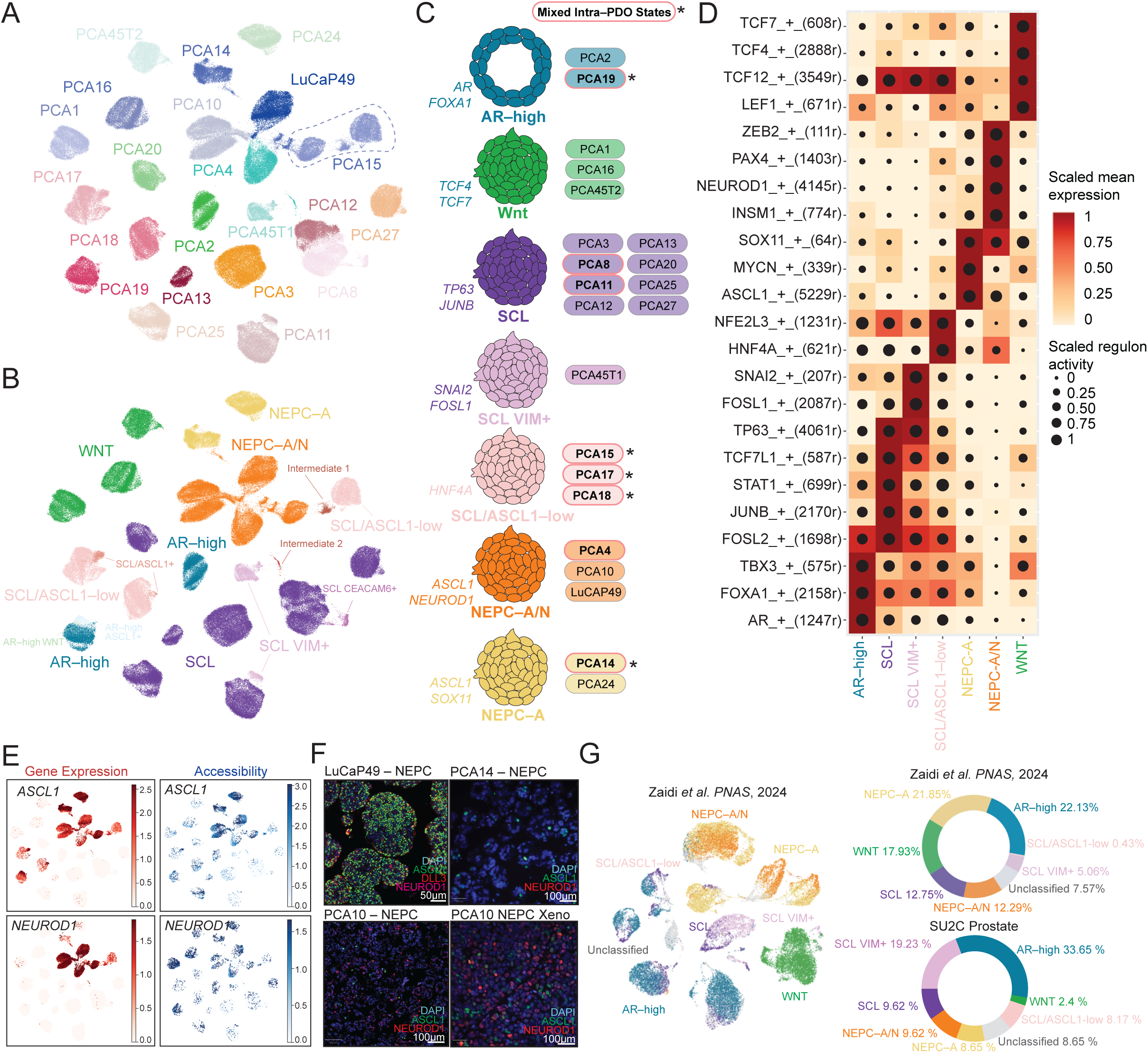
Single–cell multiome atlas of lineage states across prostate cancer organoids (PDOs). **(A)** UMAP projection of 22 PDO profiled by single–cell RNA and ATAC (sc–multiome) profiling, colored by PDO identity. Dotted line highlights PCA15 – an example of a PDO with multiple subpopulations. **(B)** UMAP colored by seven major lineage states, identifying AR–high, WNT, SCL, SCL VIM+, SCL/ASCL1–low, NEPC–A, NEPC–A/N, and multiple intermediate populations. **(C)** Schematic summarizing seven major lineage assignments, highlighting intra–PDO heterogeneity (asterisks and red bolded circle denote PDOs with mixed populations). **(D)** Scaled mean expression (color) and regulon activity (dot size) of differential lineage–defining transcription factors across subtypes using SCENIC+. **(E)** UMAP showing gene expression and chromatin accessibility for *ASCL1* and *NEUROD1*. **(F)** Immunofluorescence of neuroendocrine markers (ASCL1, NEUROD1, DAPI +/- DLL3) across representative NEPC PDOs. Scale bars, 50–100 μm. **(G)** UMAP of sc–RNA dataset from CRPC patient tumors [PMID: 38968122] labeled by PDO–defined lineage states using nearest template prediction (NTP) approach (left). Subtype composition in both sc–RNA–seq dataset and SU2C bulk tumors [PMID: 31061129] (right).

This approach resolved seven major subtypes (AR–high, SCL, SCL/VIM+, SCL/ASCL1–low, NEPC–A, NEPC–A/N, WNT) (Fig 1B and Fig 1C). These subtypes were largely independent of shared genomic alterations, except for WNT PDOs, which harbored alterations in Wnt pathway genes (Fig S1D). We also identified several minor and transitory populations (Fig 1B). To further refine the enhancer–driven gene–regulatory networks (GRNs) specific to individual PDOs (Fig S1E) and lineage subtypes (Fig 1D), we applied SCENIC+^15^ to integrate the regulatory basis of these states. These analyses identified subtype–specific regulatory programs consistent with prior functional and genetic observations and uncovered refined subclassifications within SCL and NEPC states. Consistent with the prior bulk–classification framework, the AR–high PDOs (PCA2, PCA19) showed increased *AR* and *FOXA1* activity, in line with their known sensitivity to AR pathway inhibitors (ARPI)^16^. Similarly, WNT PDOs (PCA1, PCA16, PCA45T2) showed high expression and regulon activity for *TCF7* and *TCF4*, consistent with canonical WNT signaling^10^ (Fig 1D).

Given the known plasticity of bulk–defined SCL and NEPC states^10^, we next leveraged sc–multiome profiling to resolve the heterogeneity within lineage groups. Notably, SCL organoids showed three distinct subtypes. SCL/ASCL1–low PDOs (PCA17, 18) lacked *AR* expression, showed high *HNF4A* activity, and exhibited increased *ASCL1* expression with accessibility profiles distinct from NEPC PDOs (Fig 1D). The SCL/VIM+ PDO (PCA45T1) displayed a distinct TF profile with high *FOSL1* and *SNAI2* activity and vimentin expression (Fig 1D, Fig S1C). The remaining SCL PDOs (PCA3, 8, 11, 12, 13, 20, 25, 27) showed more variable TF activity in *JUNB*, *FOSL2*, *TP63*, and *STAT1*, but were grouped as SCL based on shared expression of stem markers (*CD44* and *TROP2*) and their prior bulk–classification by Tang et al ^10^ (Figs 1C and 1D, Fig S1C and S1D).

As with SCL states, we examined whether NEPC PDOs exhibit additional stratification at the sc–multiome level. NEPC PDOs (PCA4, PCA10, PCA14, PCA24, LuCap49) expressed classical neuroendocrine markers, *CHGA* and *SYP*, but separated into two states: two NEPC–A PDOs (PCA14 and PCA24) showed ASCL1 expression and activity only, whereas three NEPC–A/N PDOs (PCA4, PCA10, and LuCap49) showed co–expression and activity of ASCL1 and NEUROD1 within the same cells, with lower levels of ASCL1 expression than NEPC–A (Fig 1E).

These sc–multiome defined segregation prompted us to assess whether inferred regulatory programs are reflected at the protein level. Immunofluorescence for neuroendocrine markers (*NCAM1, ASCL1, DLL3,* and *NEUROD1*) (Fig. 1F, Fig. S2) revealed substantial intra–model heterogeneity. In NEPC–A/N PDOs, ASCL1 and NEUROD1 protein levels varied substantially: a subpopulation of PCA10 cells co–expressed ASCL1 and NEUROD1, while LuCaP49 was almost uniformly ASCL1 positive. PCA14 (NEPC–A) was initially dominated by ASCL1+ cells with rare NEUROD1+ cells, but later passages showed an increased NEUROD1+ fraction, suggesting clonal drift (Fig. S2). Interestingly, ASCL1 and NEUROD1 protein distributions were not fully concordant with gene expression and chromatin accessibility, indicating additional regulatory complexity. Other lineage states (AR, TROP2, YAP/TAZ) showed expected marker patterns with occasional admixture in PDOs with mixed subpopulations (Fig S2).

We were motivated to test whether these 7 PDO–defined lineage states are representative of lineage states in advanced human prostate cancer. Using a nearest template prediction (NTP) approach with permutation–based significance testing, we analyzed single–cell RNA–seq data of metastatic CRPC from heavily–treated patients^2^ (N=14) and identified all major PDO–defined lineage states at the tumoral level (Fig 1G, Fig S1F). To account for potential sampling biases in this single cell cohort, we applied the same subtype signatures to bulk transcriptomic data from the much larger (N=429) Stand Up To Cancer (SU2C) CRPC cohort to estimate subtype composition^17^. Among the classified SU2C tumors, the AR–high subtype was the most frequent (>30%), followed by SCL/VIM+ (∼20%), SCL (∼10%), and NEPC–A and NEPC–A/N (∼20%), whereas SCL/ASCL1–low (∼8%) and WNT (∼2-3%) are rarer (Fig 1G, Fig S1G). Approximately ∼8-9% remain unclassified, likely reflecting lineage states not captured in the current collection of PDOs. Together, these findings establish a prostate cancer PDO resource with comprehensive single cell annotation that provides greater clarity and resolution of lineage states seen in metastatic CRPC patients.

### PDO Plasticity and Topic Modeling

While these lineage states define major transcriptional programs, our approach also sought to capture heterogeneity within each subtype and individual PDOs. Notably, several PDOs—particularly in the SCL group—lacked a dominant transcription and chromatin accessibility program. To quantify this heterogeneity, we applied the Inverse Simpson Index (ISI), a metric often used to measure ecological diversity. ISI captures either the effective number and evenness of expressed genes (ISI GEX) or accessible chromatin regions (ISI ATAC) within each PDO (Fig. S3A) or subtype while controlling for sample size (Fig 2A, Methods). Higher ISI values indicate increased regulatory diversity and lack a stable lineage commitment.

**Figure 2.**
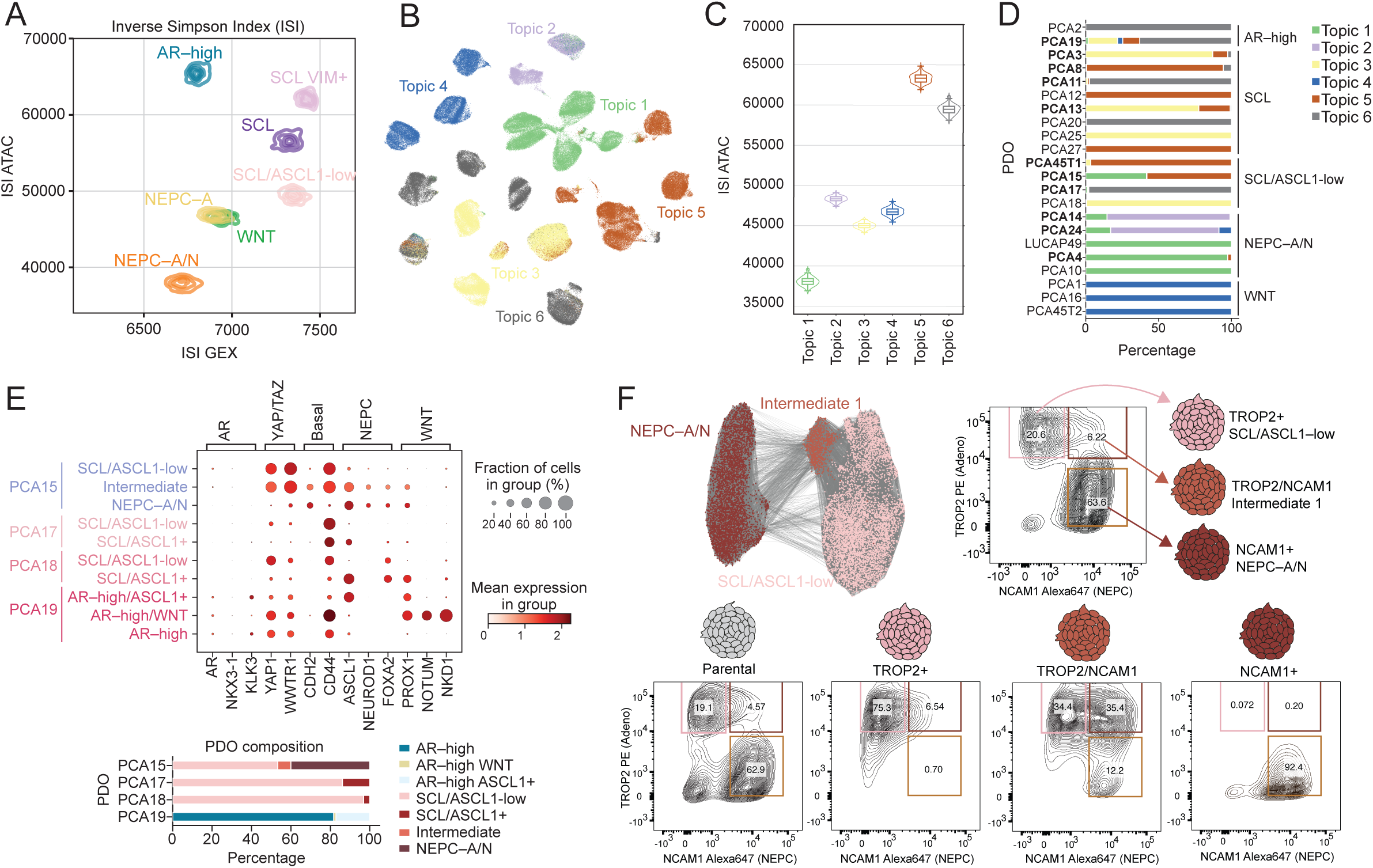
Regulatory heterogeneity and plasticity within and across prostate cancer PDOs. **(A)** Inverse Simpson Index (ISI) comparing transcriptional (GEX) and chromatin (ATAC) diversity across seven major lineage states. **(B)** Topic modeling of chromatin accessibility identifying six co–accessibility programs (Topic Modules) projected onto UMAP space. **(C)** Distribution of Inverse Simpson Index of chromatin diversity (ISI ATAC) across Topic Modules. **(D)** Fractional composition of Topic Modules within each PDOs, grouped by subtype. **(E)** Dot plot showing expression of select lineage markers across subpopulations within representative PDOs, namely PCA15, PCA17, PCA18, and PCA19 with corresponding compositional bar plots below. **(F)** UMAP showing three populations of SCL/ASCL1–low, intermediate, and NEPC–A/N found in PCA15 (left top). FACS using TROP2–PE and NCAM1–647 to identify TROP2+, NCAM1+, and double–positive intermediate populations (right). Bottom, re–plating assays of unsorted or each subpopulation, which demonstrates the lineage potential of sorted populations after 21 days of organoid culture.

SCL, SCL/ASCL1–low, and SCL/VIM+ subtypes exhibited the highest ISI GEX and ISI ATAC scores, whereas more lineage–constrained populations––including WNT, NEPC–A/N, and NEPC–A–– showed markedly lower ISI values. These ISI scores were further supported by a CRPC patient from whom two PDOs were derived independently from two different regions of the primary prostate tumor: one with high plasticity classified as SCL/VIM+ (PCA45T1) and the second with low plasticity classified as WNT(PCA45T2). This patient subsequently developed liver metastases with NEPC histology which, based on ISI, may have arisen from the SCL/VIM+ subtype. Notably, AR–high PDOs showed low ISI GEX, consistent with an androgen–dependent adenocarcinoma phenotype, but exhibited unexpectedly high ISI ATAC (Fig 2A). This discordance, exemplified by PCA19, which harbors multiple lineage populations, is consistent with its permissive chromatin landscape. Together, these findings position ISI as a potential measure of lineage plasticity that quantitatively captures population heterogeneity, supported by orthogonal biological and clinical observations.

While ISI quantifies the extent of regulatory heterogeneity, it does not resolve the underlying programs that drive diversity. To identify the chromatin accessibility profiles of these heterogenous states, we applied unbiased topic modeling [latent Dirichlet allocation (LDA)] to the chromatin accessibility data to identify co–accessibility programs (“Topic Modules”) (Fig 2B). SCL– and AR–high populations again showed considerable variability in accessibility falling into three Topic Modules (3, 5 and 6), consistent with their high plasticity. Topics 5 and 6–– enriched in AR–high and SCL states––showed the highest ISI ATAC scores (Fig 2C). Topic 5 was enriched for *FOSL1* and other *AP1* factors, whereas Topic 6 showed higher enrichment for STAT1, potentially representing previously described JAK/STAT driven states (Fig S3B).

A complementary approach that leverages multiomics data is Spectra matrix factorization on highly variable genes (“Hotspot Modules”) (Fig S3C and S3D, and GSEA analysis shown in Fig S4A). These Hotspot modules showed substantial overlap with Topic Modules, except for Topic Module 3 and Hotspot Module 5 (Fig S4B). All TFs enriched in both Topic and Hotspot modules are in Table S2 and S3, respectively. Interestingly, Hotspot and Topic Modules failed to fully separate CRPC–AR and CRPC–SCL PDOs,^8^ suggesting that some AR–high PDOs may share regulatory networks with SCL PDOs rather than being governed by fully distinct lineage–specific programs. This limitation also raises the possibility of multiple regulatory programs co–existing within individual PDOs, with implications for intra–PDO heterogeneity.

Building on this, we next asked whether regulatory programs co–exist within individual PDOs. Analysis of Topic Module composition revealed that 13 PDOs contained mixtures of two or more modules (Fig 2D), suggesting that PDO culture conditions do not select for a single dominant clone but instead maintain co–existing cellular states over time. A similar degree of subtype mixing was seen in our cluster–based annotations (Fig S4C). Focusing on these highly admixed PDOs, PCA19 (AR–high) showed three subpopulations: predominantly AR–high, with smaller WNT and ASCL1 subpopulations. PCA17 and PCA18 (SCL/ASCL1–low) both showed a dominant HNFA+ and a minor ASCL1–high (<20%) population, whereas PCA15 (mixed SCL and ASCL1) contained nearly equal SCL and NEPC (NCAM+) subpopulations with a small intermediate/hybrid subpopulation (Figs 2E and 2F).

The differential expression of the cell surface proteins TROP2 and NCAM1 within the SCL and NEPC subpopulations respectively, coupled with a high ISI score for SCL, provides an opportunity to directly address intra–tumoral plasticity through FACS–based isolation of these subpopulations. As expected, flow cytometry using TROP2–PE and NCAM2–Alexa 647 co–staining revealed three populations: TROP2–positive, NCAM1–positive, and TROP2/NCAM1–double positive. Sorting and reseeding revealed that only the double–positive subpopulation could regenerate all three lineages after 14 days, indicating a unique bipotent population that mediated NEPC transition (Fig 2F). The TROP2–positive population did give rise to a small fraction of double–positive cells (<10%), suggesting it has some potential to revert to an intermediate state whereas the NCAM1–positive cells exclusively gave rise to a homogenously NCAM1–positive population, consistent with the NEPC ISI scores predictive of low plasticity potential. Together, these results show that multiple regulatory states co–exist across and within organoids, particularly in SCL populations, and that ATAC–based measures identify organoids poised for plasticity, as exemplified by PCA15 and PCA19.

### CRISPR Screens for Synthetic Lethality

The diversity of regulatory programs within and across PDOs raises a key question of which genetic dependencies sustain specific cell states, and how intratumoral heterogeneity may influence their detection. To address this, we performed pooled CRISPR–Cas9 dropout screens in PDOs using single guide (sgRNA) containing customized libraries (Fig 3A). Unlike conventional 2D cell line screens, CRISPR screening in 3D organoid systems presents additional challenges, including maintaining library representation and achieving sufficient sensitivity to detect fitness effects in clonally expanding structures^18^. As CRISPR screens in 3D human PDO cultures have yet to be performed extensively, we first optimized key experimental parameters to ensure robust and interpretable gene depletion results (Methods, Fig S5A–B). For this, we generated mini–CRISPR libraries targeting 843 differentially expressed genes across PDO subtypes and stratified them into three functional pools: Pool 1–TF (transcription factors), Pool 2–Epigenetics/Kinases (epigenetic regulators and kinases), and Pool 3–Druggable (druggable targets). Each pool contained four sgRNA for 281 differentially expressed genes with 5% non–targeting and SAFE control guides (Methods, Table S4).

**Figure 3.**
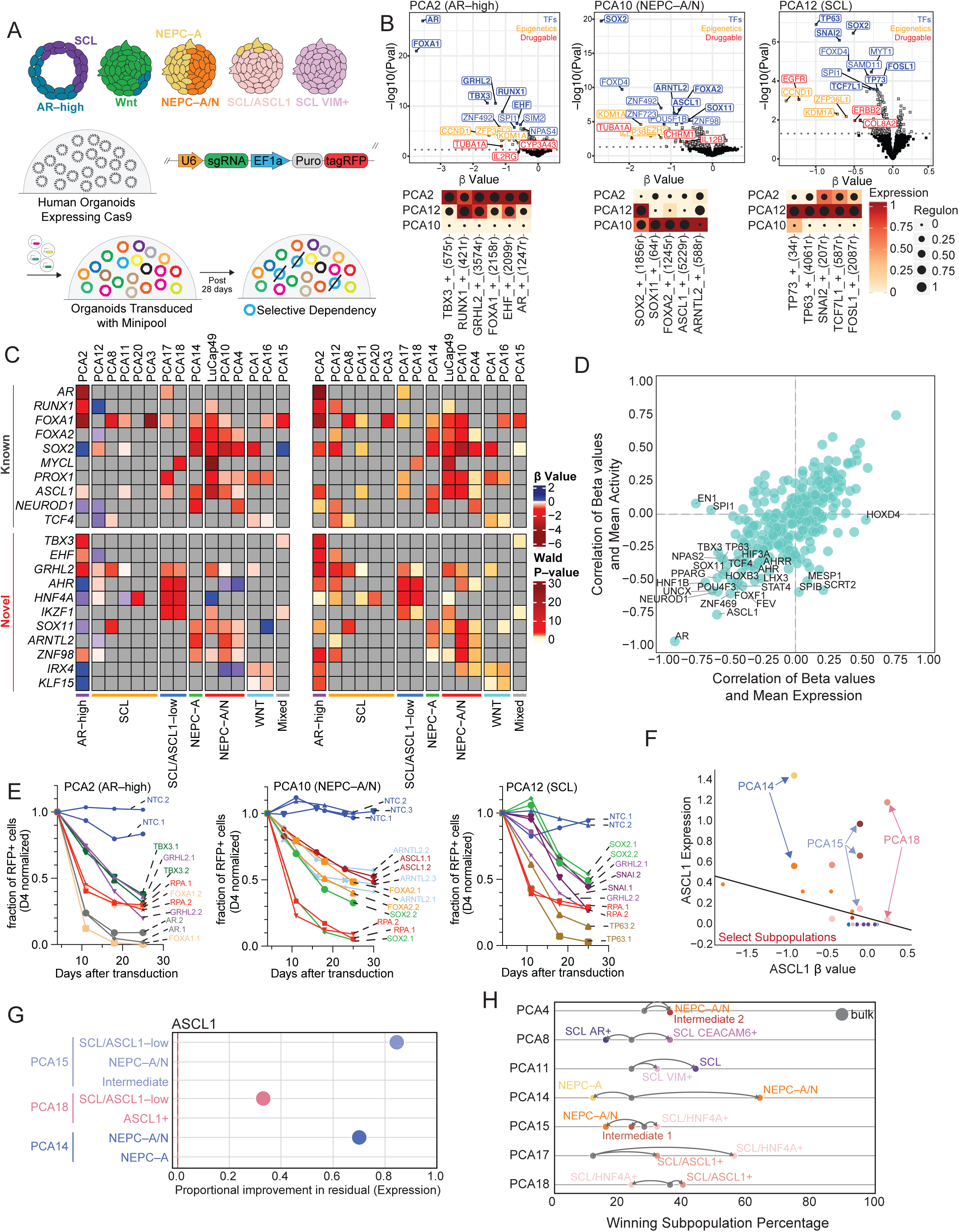
Lineage–resolved CRISPR screening identifies subtype–specific dependencies. **(A)** Schematic of pooled CRISPR–Cas9 screening in PDOs of different lineages. **(B)** Volcano plots showing gene–level depletion (β value) and significance (–log10 P value) for representative AR–high (PCA2), NEPC–A/N (PCA10), and SCL (PCA12) PDOs, with selected top 10 transcription factor, and top 3 epigenetic and druggable targets highlighted. Bolded TFs show overlap with SCENIC+ analysis for enrichment in particular PDO and shown as a heatmap below with scaled mean expression (color) and regulon activity (dot size) using SCENIC+. **(C)** Heatmaps of CRISPR dependencies across PDOs for known (top) and newly identified (bottom) transcription factor dependencies with left showing depletion value (β value) and right showing significance (–log10 P value). Only significant depleters are shown (P< 0.01). **(D)** Correlation between CRISPR depletion (β values) and gene expression or regulon activity, highlighting candidate dependencies. **(E)** sgRNA competition assays validating lineage–specific dependencies across PDOs over time with fraction of RFP shown on y–axis. Data is normalized to day 4 of transduction efficiency. **(F)** Subpopulation–resolved analysis of *ASCL1* dependency showing divergence between bulk and subpopulation–specific effects, namely PCA14, PCA15, and PCA18. **(G)** Improvement in model residuals when incorporating subpopulation–level expression for *ASCL1* across select PDOs. **(H)** Distribution of “winning” subpopulations that best explain gene–level depletion signals across TFs screens in multiple admixed PDOs.

To establish feasibility and robustness, we utilized three PDOs with divergent lineage programs and chromatin states that may pose different technical challenges: AR–high (PCA2), NEPC–A/N (PCA10), and SCL (PCA12). PDOs stably expressing Cas9 were transduced with each library, and genomic DNA was collected at baseline (T_0_) and after 21 or 28 days (T_1_ or T_2_, ∼8–10 doublings). Gene level dependencies were quantified using MAGeCK based on sgRNA depletion (β value) and statistical significance (Wald P value) (Table S5). Consistent with lineage–specific dependencies for Pool 1–TF screens, the AR–high PDO (PCA2) showed strong depletion for *AR* and *FOXA1*, the NEPC–A/N PDO (PCA10) showed depletion for *SOX2*, *ASCL1* and *FOXA2*, and the SCL PDO (PCA12) was dependent on *TP63* and *SNAI2*. Notably, the top subtype–specific TF dependencies corresponded to the TF expression and regulon activity (SCENIC+) in the sc–multiome atlas (Fig 3B, Table 1, Table S5).

**TABLE 1.**

| **PDO** | **Gene** | **Beta value** | **Wald P value** | **Subtype** |
| --- | --- | --- | --- | --- |
| LuCap49 | *MYCL* | -4.306 | 3.71x10<sup>-16</sup> | NEPC-A/N |
| PCA2 | *FOXA1* | -3.745 | 1.07x10<sup>-21</sup> | AR-high |
| PCA2 | *AR* | -3.549 | 1.06x10<sup>-27</sup> | AR-high |
| LuCap49 | *ASCL1* | -2.797 | 3.99x10<sup>-07</sup> | NEPC-A/N |
| PCA10 | *SOX2* | -2.673 | 1.91x10<sup>-20</sup> | NEPC-A/N |
| PCA14 | *SOX2* | -2.326 | 6.52x10<sup>-06</sup> | NEPC-A |
| PCA17 | *HNF4A* | -2.033 | 6.57x10<sup>-05</sup> | SCL/ASCL1-low |
| PCA18 | *AHR* | -1.948 | 1.63x10<sup>-05</sup> | SCL/ASCL1-low |
| PCA18 | *HNF4A* | -1.868 | 7.74x10<sup>-05</sup> | SCL/ASCL1-low |
| PCA2 | *TBX3* | -1.562 | 2.56x10<sup>-11</sup> | AR-high |
| PCA17 | *AHR* | -1.559 | 5.67x10<sup>-05</sup> | SCL/ASCL1-low |
| PCA4 | *SOX2* | -1.348 | 1.78x10<sup>-06</sup> | NEPC-A/N |
| PCA14 | *FOXA2* | -1.332 | 1.60x10<sup>-03</sup> | NEPC-A |
| PCA2 | *GRHL2* | -1.308 | 2.91x10<sup>-11</sup> | AR-high |
| LuCap49 | *SOX2* | -1.256 | 1.64x10<sup>-04</sup> | NEPC-A/N |
| LuCap49 | *PROX1* | -1.156 | 3.52x10<sup>-04</sup> | NEPC-A/N |
| PCA2 | *RUNX1* | -1.098 | 1.44x10<sup>-09</sup> | AR-high |
| PCA18 | *IKZF1* | -1.042 | 3.16x10<sup>-02</sup> | SCL/ASCL1-low |
| PCA17 | *IKZF1* | -1.034 | 1.29x10<sup>-03</sup> | SCL/ASCL1-low |
| LuCap49 | *FOXA2* | -1.011 | 2.10x10<sup>-03</sup> | NEPC-A/N |
| PCA12 | *TP63* | -0.994 | 1.27x10<sup>-07</sup> | SCL |
| PCA14 | *ARNTL2* | -0.974 | 8.92x10<sup>-03</sup> | NEPC-A |
| PCA10 | *ARNTL2* | -0.911 | 7.48x10<sup>-07</sup> | NEPC-A/N |
| PCA14 | *ASCL1* | -0.909 | 2.75x10<sup>-03</sup> | NEPC-A |
| PCA10 | *FOXA2* | -0.856 | 8.59x10<sup>-07</sup> | NEPC-A/N |
| PCA1 | *PROX1* | -0.815 | 1.36x10<sup>-03</sup> | WNT |
| PCA10 | *ASCL1* | -0.802 | 1.80x10<sup>-07</sup> | NEPC-A/N |
| PCA10 | *ZNF98* | -0.715 | 1.01x10<sup>-05</sup> | NEPC-A/N |
| PCA16 | *PROX1* | -0.715 | 1.23x10<sup>-02</sup> | WNT |
| PCA12 | *SNAI2* | -0.693 | 8.09x10<sup>-07</sup> | SCL |
| PCA14 | *ZNF98* | -0.641 | 4.72x10<sup>-02</sup> | NEPC-A |
| PCA1 | *TCF7L1* | -0.494 | 3.02x10<sup>-02</sup> | WNT |
| PCA4 | *FOXA2* | -0.431 | 3.85x10<sup>-02</sup> | NEPC-A/N |
| PCA1 | *IRX4* | -0.417 | 2.22x10<sup>-02</sup> | WNT |
| PCA16 | *IRX4* | -0.415 | 1.39x10<sup>-02</sup> | WNT |
| PCA4 | *ARNTL2* | -0.375 | 1.49x10<sup>-02</sup> | NEPC-A/N |
| PCA16 | *KLF15* | -0.344 | 2.09x10<sup>-02</sup> | WNT |
| PCA1 | *TCF4* | -0.290 | 1.76x10<sup>-02</sup> | WNT |
| LuCap49 | *ZNF98* | -0.259 | 3.84x10<sup>-02</sup> | NEPC-A/N |
| PCA4 | *ASCL1* | -0.186 | 3.32x10<sup>-02</sup> | NEPC-A/N |
| PCA4 | *ZNF98* | -0.179 | 1.91x10<sup>-02</sup> | NEPC-A/N |
| PCA16 | *TCF4* | -0.111 | 3.68x10<sup>-02</sup> | WNT |

Figure legend: AR, androgen-receptor; NEPC, neuroendocrine prostate cancer; PDO, patient-derived organoid; SCL, stem-cell-like; WNT, WNT signaling.

Pool 2–Epigenetics/Kinases and Pool 3–Druggable screens also identified subtype–specific (as well as shared) vulnerabilities. *LSD1/KDM1A* and *TUBA1A* were depleted across all three PDOs with the strongest depletion in the NEPC–A/N PDO (PCA10). *ZFP36L2* and *EZH2* were also depleted strongly in PCA10 but more modestly in the AR–high PDO (PCA2), consistent with known function in these lineage states^13, 19, 20^. Several genes stood out for their strong and unique depletion including *HNRNPCL1*, *PTHLH*, *COL4A1*, and *IL12B* in the NEPC–A/N PDO. *HKDC1* (a hexokinase) and *IL2RG* were uniquely depleted in the AR–high PDO, whereas *ZFP36L1* and *EGFR* were uniquely depleted in the SCL PDO (Fig 3B, Table S5). Overall, the mini–CRISPR screen confirmed expected lineage–specific dependencies while revealing additional subtype-specific vulnerabilities.

Given our successful optimization of small scale CRISPR screens––defined by the robust depletion of known TFs in distinct lineage states––we expanded the Pool 1–TF screen to an additional 12 PDOs with the goal of constructing a TF dependency atlas across prostate cancer lineage states. All four NEPC PDOs showed significant *ASCL1* dependence, as expected. LuCaP49 showed the strongest *ASCL1* dependence, along with dependencies on *MYCL* and *PROX1*, whereas PCA4 showed greater dependence on *NEUROD1*, together with ASCL1 dependence. Other pan–NEPC TF dependencies include *SOX2*, *FOXA1* and *FOXA2* with higher β–values overall for *FOXA2* versus *FOXA1*, consistent with the known shift from *FOXA1* to *FOXA2* expression when CRPC transitions to NEPC (Fig 3C, Table 1, Table S5).

Beyond canonical NEPC regulators, we identified three previously unreported NEPC–selective dependencies. *SOX11* sgRNAs were depleted in all NEPC PDOs. sgRNAs for *ARNTL2* (aryl hydrocarbon receptor nuclear translocator like 2), a circadian rhythm regulator that dimerizes to CLOCK protein and is implicated in advanced lung adenocarcinoma^21^, were depleted in 3 of 4 NEPC PDOs but not in other PDO subgroups. Lastly, sgRNAs for the zinc finger TF *ZNF98* were depleted across all 4 NEPC PDOs (Fig 3C, Table 1, Table S5).

In the AR–high state (PCA2), top ranking dependencies were *TBX3*, *GRHL2*, *RUNX1*, and *EHF*, in addition to well established dependencies on *AR* and *FOXA1* (Fig 3C, Table 1, Table S5). For WNT subtype PDOs (PCA1, 16), the screen revealed dependencies on *PROX1*, *IRX4* and *KLF15*, as well as known dependencies on *TCF7L1* and *TCF4* (Fig 3C, Table 1, Table S5). For SCL/ASCL1–low subtype PDOs (PCA17, 18), we identified three novel TF candidates: *AHR*, *HNF4A*, and *IKZF1*. We did not identity any consistent SCL subtype dependencies, in line with the high level of heterogeneity (ISI) seen in our sc–multiomics analyses (Fig 3C, Table 1, Table S5).

To address whether pooled functional screens could be scaled beyond the size of our mini–pooled libraries, we next screened a comprehensive TF and epigenetics factors sgRNA library comprising of 1,925 genes (Table S6) on three representative PDO models: PCA2 (AR–high), PCA12 (SCL), and PCA10 (NEPC–A/N). We first noted that depletion profiles for significantly depleted genes were highly concordant with those generated using the mini-Pool 1–TF and mini–Pool 2–Epigenetic/Kinases libraries, demonstrating reproducibility of β–values across screening platforms and confirming that library complexity can be scaled without introducing bottlenecking artifacts (Fig S5C). As expected, core essential genes, as defined by DepMap and Hart *et al* ^22^, were strongly depleted. Beyond these shared essentialities, the expanded library revealed subtype-specific dependencies not captured in the smaller pools, including *E2F3*, *TOX4*, and *SOX4* in NEPC–A/N. The *E2F3* dependency is notable because PCA10 is *RB1*–null, consistent with reduced *CCND1* dependence and increased reliance on E2F–mediated proliferation. Other top ranked dependencies include *KAT7* and *TTF1* in SCL PCA12, as well as *HOXA13,* a known regulator of luminal identity, and *ACO2* in AR–high PCA2 (Fig S5D, Table S7).

We next asked whether elevated gene expression and activity predict dependency, focusing on “dosage–sensitive” genes whose dependency is highly correlated with expression/activity, as these represent high–priority candidates for functional validation. By integrating CRISPR dependency data with TF expression and/or accessibility, we identified 23 genes with moderate to strong (anti–) correlation between expression or activity and depletion [abs(r) ≥ 0.5]. In addition to genes such as *AR* and *ASCL1*, we identified *HNF1B*, *SOX11*, *UNCX*, and *NPAS2* (Fig 3D, Table S8) as novel candidates. To validate subtype–specific and gene–dose–specific dependencies revealed from the CRISPR screens, we performed competition assays. In all cases, sgRNA competition assays recapitulated the magnitude and direction of depletion observed in pooled screens, further confirming the robustness and specificity of the identified dependencies (Fig 3E, Fig S5E, Table S9). Notably, several of these dependencies exhibited depletion effects comparable to or exceeding those observed for the canonical common essential gene *RPA3*, underscoring their functional significance. Together, these data provide a lineage–resolved map of functional dependencies across prostate cancer states, showing that while lineage identity strongly shapes functional dependencies, poorly defined states such as SCL PDOs exhibit heterogeneous and less consistent vulnerabilities.

### Effects of Intra–PDO Structure and CRISPR Depletion

Our integrated datasets further enabled us to directly assess how intra–PDO heterogeneity shapes gene–level depletion signals in pooled CRISPR screens. As depletion in heterogenous PDOs reflects the aggregate contribution of multiple subpopulations rather than a single uniform dependency, we implemented a leave–one–out modeling framework in which each PDO was excluded during model fitting, and gene expression or regulatory activity was inferred from CRISPR depletion (β values). Model performance was evaluated using the proportion of variance explained (R²), enabling direct comparison of bulk *versus* subpopulation–resolved predictions (Fig S6A).

*ASCL1* served as a prime example of this effect. Despite high expression across multiple NEPC PDOs, its depletion varied widely. At the bulk level, *ASCL1* expression and depletion were weakly correlated (Fig S6B). However, subpopulation–resolved analysis revealed that depletion was in fact driven by specific NEPC subpopulations (Fig 3F and Fig S6C). In NEPC PDO PCA14, replacing bulk averages with subpopulation–specific values showed that *ASCL1* depletion was primarily attributable to a NEPC–A population lacking *NEUROD1*, resulting in a substantial proportional improvement in residuals and overall model fit (Fig 3G). Similar patterns with *ASCL1* were observed in PCA15 and PCA18 (Fig 3G, Fig S6D). More broadly across other TFs, two distinct patterns were observed in PDOs where a single subpopulation accounted for most of the depletion signals (e.g. PCA14), while other depletion signals were distributed across multiple subpopulations (e.g. PCA15) (Figs S6D–G). By incorporating subpopulation–resolved modeling, R² values for predicting key TF candidates improved compared to bulk estimates (Figs S6H). To quantify the influence of subpopulation dynamics in each PDO, we calculated the frequency with which each subpopulation emerged as the best–performing predictor (“winning subpopulation”) of gene–level depletion across all TFs included in our screens (Fig. 3H). The improved fit of subpopulation–resolved models over bulk estimates suggests that population vulnerability to TF depletion is strongly influenced by its underlying subpopulation architecture.

We then evaluated whether the expression of a given TF within a specific subpopulation determined which subpopulation predominantly drove the overall depletion. For each PDO, we applied a G–test of independence to assess whether the subpopulation providing the greatest improvement in model fit for a given TF was statistically independent of the subpopulation exhibiting the lowest expression of that TF. Low–expressing populations were disproportionately represented among the winning clones, indicating that they generally drive aggregate depletion (Table S10). Together, these results show that gene–level dependencies in heterogeneous PDOs are shaped by specific subpopulations, with resistant or low–expression populations disproportionately influencing CRISPR depletion outcomes, providing a framework to resolve lineage–specific vulnerabilities within complex PDO systems.

### AHR as a target for SCL and SCL/ASLC1 PDOs

Our sub–population–resolved framework revealed identification of lineage–restricted dependencies that are partially obscured in bulk analyses. Among these, AHR emerged as a top candidate selective vulnerability in SCL/ASCL1–low PDOs (PCA17 and PCA18) in our pooled screens (Fig 4A), with a more modest effect observed in the SCL organoid PCA12 (refer to Fig 3C). We therefore sought to determine whether AHR represents a functional and pharmacologically actionable target across PDO subtypes.

**Figure 4.**
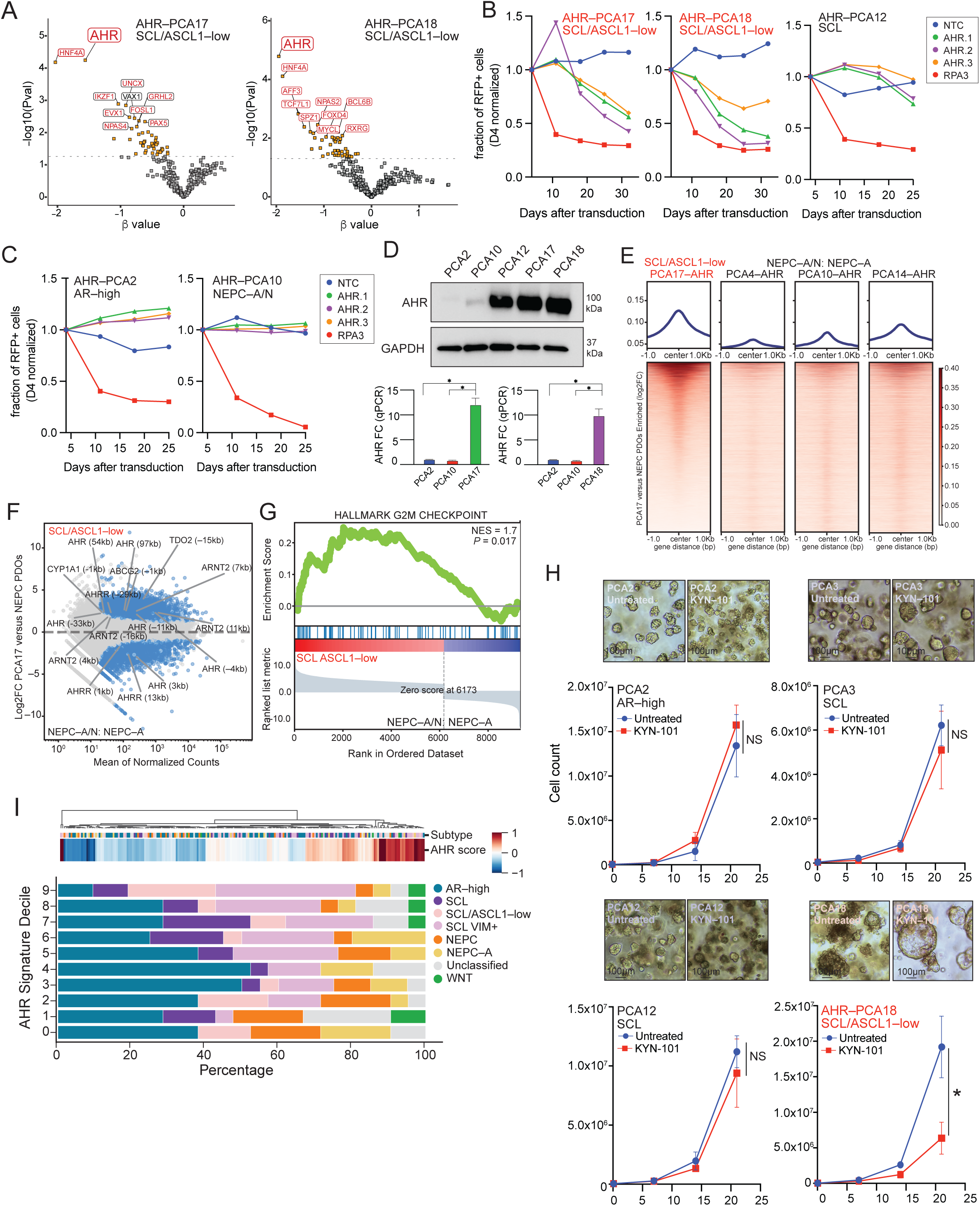
AHR is a lineage–specific dependency in SCL/ASCL1–low and SCL/VIM+ prostate cancer. **(A)** Volcano plots showing AHR as a selectively depleted gene in SCL/ASCL1–low PDOs (PCA17, PCA18). **(B)** sgRNA competition assays validating AHR dependency in SCL/ASCL1–low, PCA17 and PCA18, and more modestly in SCL PDOs, PCA12. Shown is fraction of RFP–positive cells, normalized to D4 (transduction efficiency). **(C)** AHR knockout effects across AR–high and NEPC PDOs showing increased growth or non–effect, respectively. Shown is fraction of RFP–positive cells, normalized to D4 (transduction efficiency). **(D)** Immunoblot and qPCR analysis confirming elevated AHR expression in SCL/ASCL1–low PDOs. **(E)** Volcano plot showing AHR ChIP–seq signal increased in SCL/ASCL1–low PCA17 *versus* NEPC PDOs PCA4, 10, and 14. **(F)** Differential expression of AHR target genes when analyzing AHR CHIP–seq comparing SCL/ASCL1–low and NEPC states. **(G)** Gene set enrichment analysis showing enrichment of G2/M checkpoint pathways in SCL/ASCL1–low PDOs *versus* NEPC. **(H)** Pharmacologic inhibition of AHR (KYN–101 at 1µM) across PDOs showing selective growth suppression in SCL/ASCL1–low model PCA18 with morphologic changes, but not AR–high PDOs. **(I)** Distribution of AHR signature scores across SU2C prostate cancer samples, stratified by lineage subtype showing enrichment of specific SCL subgroups in top decile of AHR scores.

Using three independent AHR sgRNAs, we validated AHR dependency in competition assays in SCL/ASCL1–low subtypes PDO (PCA17, 18), observing depletion by ∼17–18 days, with a more modest effect in SCL subtype PDO PCA12 (Fig 4B). In contrast, AHR knockout conferred a proliferative advantage in the AR–high PDO PCA2 but had no effect in NEPC–A/N subtype PDO PCA10 (Fig 4C) ––findings supporting a context–dependent role for AHR across lineage states.

To determine whether these differential dependencies reflect lineage–specific pathway activity, we interrogated AHR signaling across the multiome atlas. AHR expression was highest in SCL/ASCL1–low, SCL–VIM+, and SCL subtype PDOs, together with CYP1B1, a canonical AHR target gene, whereas expression of its obligate dimerization partner ARNT was relatively uniform across states (Fig S7A). These findings were independently validated by qPCR and immunoblotting, confirming that SCL/ASCL1–low PDOs express high levels of AHR (mRNA and protein) relative to AR–high and NEPC PDOs (Fig 4D).

Having confirmed differential AHR expression, we next asked whether this correlated with chromatin occupancy. AHR ChIP–seq revealed increased DNA binding in the SCL/ASCL1–low PDO PCA17 compared to three NEPC subtype PDOs (PCA10, PCA14, PCA4) (Fig 4E, Fig S7B and S7C). Furthermore, the AHR peaks in PCA17 mapped to canonical AHR target genes (*AHRR*, *CYP1A1*, *CYP2S1*, UGT1A family members, and *ABCG2*) and, through GSEA analysis, showed enrichment of AHR signaling and, importantly, G2/M checkpoint pathways (Fig 4F–G) (Table S11 and S12).

Having implicated AHR as a lineage–specific dependency within a CRPC SCL subtype, we next asked whether pharmacological inhibition of AHR could reproduce the dependency seen with CRISPR depletion. To explore this possibility, we treated 4 PDOs representing different lineage states with KYN–101, a small molecule antagonist that blocks ligand–induced AHR activation. As expected, expression of the target gene CYP1A1 was reduced following KYN–101 treatment (1µM) (Fig S7D). Consistent with the CRISPR data, KYN–101 treatment significantly impaired growth in the SCL/ASCL1–low PDO PCA18 but not in the AR–high PDO PCA2 (of note, as with the CRISPR knockout, PCA2 cells treated with KYN-101 showed increased growth, *albeit* not statistically significant) (Fig. 4H, Fig S7E).

To define the clinical context in which AHR–high prostate tumors might be targeted, we queried RNA–seq data from the SU2C cohort and identified a subset of CRPC patients with high AHR signature scores (Table S13). Furthermore, and strikingly, the top decile of AHR signature–positive patients were enriched for SCL/ASCL1–low or SCL/VIM+ lineage states, suggesting that AHR activity supports maintenance of these SCL lineages (Fig. 4I). Collectively, these findings demonstrate that AHR activity is preferentially enriched in a subset of SCL states, where it supports lineage maintenance and cellular fitness, highlighting the importance of cellular context in determining both genetic dependency and therapeutic response.

## DISCUSSION

Tumor heterogeneity and lineage plasticity represent fundamental barriers to durable therapeutic control broadly across advanced solid tumors. While transcriptional and epigenetic profiling has increasingly resolved the diversity of tumor cell states, translating these descriptive atlases into actionable therapeutic strategies has remained challenging^2, 23, 24^. A major obstacle has been the lack of experimental systems that simultaneously preserve patient–relevant heterogeneity and enable scalable functional interrogation. Prostate cancer, in particular, remains underrepresented in large–scale functional genomics resources such as DepMap. Here, we address this gap by integrating single–cell multiomics profiling with pooled CRISPR–Cas9 screening across a large panel of prostate cancer PDOs to link cell state identity, regulatory programs, and functional dependencies within heterogenous human tumor models. Together, this integrated PDO atlas and screening platform provides a scalable, patient–relevant resource to define functional dependencies in advanced prostate cancer (Fig. 5).

**Figure 5.**
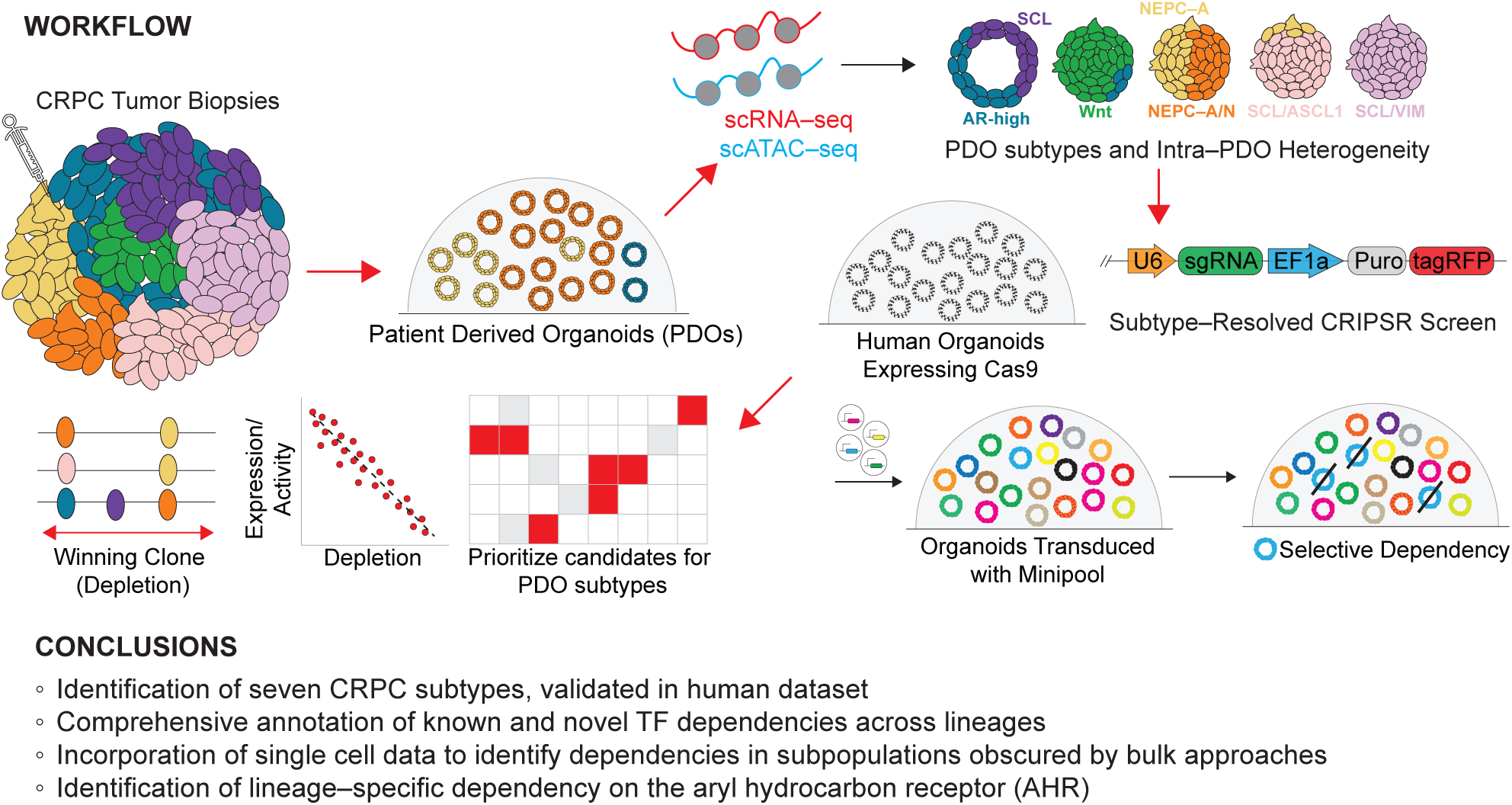
Workflow and major conclusions from integrated single–cell multiomics and CRISPR screening of heterogeneous, lineage plastic patient–derived prostate cancer organoids.

A central conceptual advance of our work is the demonstration that tumoral heterogeneity fundamentally shapes the interpretation and predictive power of pooled CRISPR screens. Conventional functional genomics screens have largely relied on relatively homogeneous cancer cell lines representative of different lineages, which implicitly assume uniform dependency landscapes^5, 25^. Our results reveal that screens performed in PDOs, which capture the heterogeneity typically seen in patients, produce gene–level depletion signals reflect the composite of multiple subpopulations with distinct transcriptional and chromatin states. Critically, we find that the subpopulation with the lowest dependency on a given gene—the "limiting subpopulation"—disproportionately determines aggregate depletion signals, presumably because this resistant population can expand and mask vulnerabilities present in other subpopulations.

This principle has two important corollaries. First, strong lineage–restricted dependencies may appear attenuated—or even undetectable—when averaged across mixed populations, as exemplified by PCA15, where extensive admixture of SCL, neuroendocrine, and intermediate states resulted in minimal depletion of most lineage–specific regulators. Second, robust depletion of genes with low mean expression can reflect vulnerabilities confined to a functionally critical subpopulation that nonetheless limits overall population fitness; strong depletion of *HNF4A* and *AHR* in PCA17, for instance, was driven by a restricted SCL/ASCL1–low population despite low bulk expression. These observations provide a mechanistic explanation for the poor predictive performance often observed when dependencies identified in cell lines are extrapolated to patient tumors and suggest that therapeutic target selection should prioritize the activity of relevant cell states rather than population averages.

Beyond resolving technical limitations of pooled screening, our work provides new biological insight into the structure of prostate cancer plasticity. By constructing a single cell multiome atlas spanning 22 PDOs, we delineated seven major lineage states. Most notable was the refined subtyping of the previously bulk–seq–defined SCL subgroup (through bulk analyses), from which we defined three distinct cell states: SCL, SCL/ASCL1–low, and SCL/VIM+, along with the heterogeneity of NEPC (NEPC–A, NEPC–A/N), pertaining to ASCL1 or NEUROD1 expression and activity^26, 27^. Importantly, we identified intermediate and hybrid populations––often masked in bulk analyses––that exhibited a high level of diversity in chromatin accessibility and transcriptional plasticity. Quantitative measures of heterogeneity, including inverse Simpson indices and topic modeling, revealed that SCL and related AR–low states exhibit the greatest regulatory variability, consistent with their proposed role as reservoirs of lineage plasticity^2, 26, 28^.

Functional CRISPR screening across these states uncovered both shared and PDO–specific dependencies, validating known regulators while also revealing previously unrecognized vulnerabilities. Established lineage regulators, such as *AR* and *FOXA1* in AR–high, *TCF4* in WNT, and *ASCL1* and *SOX2* in NEPC^10^, emerged as top dependencies in their respective subtypes, providing internal validation of our screening approach in 3D PDO cultures. Additional subtype–specific dependences, such as *TBX3* and *EHF* in AR–high, *HNF4A* and *AHR* in SCL/ASCL1–low, and *SOX11* and *ARNTL2* in NEPC warrant further investigation. Moreover, the variable dependency of *ASCL1* and *NEUROD1* in different NEPC PDOs suggests that subpopulation structure impacts the effect of gene depletion at the time of screening.

Among these findings, the identification of AHR as a selective dependency in the SCL/ASCL1–low state offers potential for near term therapeutic relevance. AHR has been studied primarily in the context of xenobiotic metabolism and immune regulation, with limited evidence linking it to specific prostate cancer lineage states^29, 30^. Our integrated genetic, pharmacologic, and epigenomic analyses show that AHR activity is enriched in SCL/ASCL1–low population, where it supports proliferation and lineage maintenance. Notably, this dependency is highly context–specific as AHR inhibition promotes growth in AR–high tumors, underscoring the importance of lineage state in determining therapeutic response if/when AHR–directed therapy is explored clinically. This SCL/ASCL1–low state is not well represented in existing DepMap prostate cancer cell lines, highlighting a key limitation of current *in vitro* models and emphasizing the value of PDO systems for capturing clinically relevant, yet underrepresented, lineage states.

While our study establishes a scalable and integrative framework, several limitations warrant consideration. First, although CRISPR screening in PDOs can be extended to large gene libraries, genome–scale screens remain constrained by the dominance of core essential genes in pooled depletion readouts, which can obscure context–specific dependencies. Our PDO panel also has limited representation of AR–high prostate cancer states, reflecting challenges in establishing and maintaining these models and potentially biasing dependencies toward AR–low and lineage-plastic phenotypes. Second, while PDOs preserve tumor cell–intrinsic heterogeneity, they lack immune and stromal components that influence lineage stability and therapeutic response *in vivo*. Finally, organoid passaging may alter clonal composition. Although drift was observed in specific cases (e.g., PCA14), most PDOs retained diverse lineage states, supporting preservation of patient–relevant heterogeneity.

Finally, these findings are likely to extend beyond prostate cancer, as lineage plasticity and cellular heterogeneity are common across solid tumors. Integrating single–cell multiomics with functional genomics in patient–derived models provide a general framework to resolve state–specific vulnerabilities and interpret screening results in heterogeneous systems. Accounting for heterogeneity will be essential for translating functional genomics into effective therapeutic strategies.

## Acknowledgements

We thank the members of the Sawyers, Zaidi, and Calico Life Sciences teams for their helpful critiques and vibrant discussions. We also thank the Feng lab for critical discussions regarding the role of AHR in prostate cancer. We are grateful for the support and assistance of the Epigenetics Research Innovation Lab, the Single Cell Analytics Innovation Lab, and the Molecular Cytology Core. We also acknowledge the use of the Integrated Genomics Operation Core, funded by the NCI Cancer Center Support Grant (CCSG, P30 CA08748), Cycle for Survival, and the Marie-Josée and Henry R. Kravis Center for Molecular Oncology. S.Z. is supported by the Prostate Cancer Foundation TACTICAL Award, the Burroughs Wellcome Fund Career Award for Medical Scientists, and NCI K08CA282978. C.L.S. was supported by the Howard Hughes Medical Institute, Calico Life Sciences LLC, and NIH grants CA193837, CA092629, CA265768, and CA008748.

## Author Contributions

S.Z. and C.L.S. conceived this study. S.Z. and C.L.S. designed the experiments and interpreted the results. M.B., A.F., and Z.C. performed and executed computational work. S.Z., C.B., T.P.P., and D.H.K. performed CRISPR screens and wet–lab experimentation. S.Z., D.H.K., C.K.W., and Y.C. generated and cultured organoids. R.C. oversaw library preparation and sequencing of single–cell multiome data. S.N., S.Z., and S.M. designed and generated CRISPR libraries. P.J.H. (and core) generated ChIP–seq data. W.K. performed immunofluorescence studies. H.J., J.C., and S.M. analyzed CRISPR data. S.Z., C.B., D.Z., and W.F. designed and executed AHR experiments. S.Z., M.B., Z.C., A.F., and C.L.S. wrote the manuscript.

## METHODS

### SINGLE–CELL MULTIOME DATA

#### Data pre–processing

The FASTQ files of the single–cell multiome data were processed by sample with cellranger–arc *v.* 2.0.0, except for samples PCA13, PCA19, PCA23, and PCA27, which were processed with *v.* 2.0.2. FASTQs were aligned against the reference genome hg38. All samples were aggregated using the cellranger–arc aggr function (*v.* 2.0.2). Genomic regions of the peaks were annotated with the HOMER^1^ annotatePeaks.pl function, *v.* 4.11, using the cellranger-arc refdata–cellranger–arc–GRCh38-2020–A–2.0.0 genes.gtf file.

#### Quality control

Cell filtering criteria were computed *per* sample and separately *per* modality, and cells were then assessed independently for the quality of gene expression and chromatin accessibility data. To address sample–to–sample variability in these metrics, we set two types of QC thresholds for each metric: permissive global thresholds and sample–wise percentile thresholds. For each metric, the 2nd and 98th percentiles were computed *per* sample as sample–wise cutoffs. An exception was the fraction of mitochondrial reads, for which the 2nd percentile cutoff was not used, as low fractions of mitochondrial reads do not present a disadvantage. A cell was deemed to pass QC in a modality if it failed no more than one of the tested QC criteria. Cells were retained when both modalities passed QC.

For single–cell gene expression (GEX) data, the number of total counts *per* cell, number of genes expressed *per* cell, and fraction of mitochondrial counts *per* cell were used as QC metrics. The following permissive global thresholds were applied in addition to the sample–wise percentile thresholds described above: library size between 2,500 and 20,000 UMIs, number of expressed genes between 200 and 10,000, and fraction of mitochondrial reads below 40%. After initial filtering, doublets were detected using the scrublet^2^ approach (*v.* 0.2.3) for gene expression data with parameters min_counts = 2, min_cells = 3, vscore_percentile = 85, n_pc = 50, expected_doublet_rate = 0.02, sim_doublet_ratio = 3, and n_neighbors = 15; cells with doublet_score > 0.14 were removed.

For single–cell chromatin (ATAC) accessibility data, the number of total counts *per* cell and number of peaks detected *per* cell were used as QC metrics. The following permissive global thresholds were used in addition to the sample–wise percentile thresholds described above: library size between 1,000 and 40,000 UMIs and number of detected peaks per cell between 500 and 20,000. In addition, we added QC criteria for fraction of reads in peaks (FrIP score), TSS enrichment score, and log unique fragments. The pycistopic^3^ package *v.* 1.10.12 in Python 3.8.17 was used to compute these additional metrics, and we added three additional QC criteria: FrIP scores had to be greater than 0.4, except for PCA17, where the threshold was 0.35; TSS enrichment had to be above 4; and log unique fragments had to pass a manually curated per-sample threshold. The resulting dataset consisted of 195,180 cells with multimodal profiles. Genes were filtered by computing the number of cells expressing a gene and filtering out all genes that were expressed in fewer than 3 cells. To filter peaks, a threshold of 50 cells per peak was used.

#### Normalization and feature selection

For single–cell GEX, normalization was performed by a shifted log–transformation of size-scaled counts^4^ using the scanpy functions pp.normalize_total(target_sum = 1e4) followed by pp.log1p. For single–cell ATAC, an analogous CPM–style log–normalization was stored as the norm layer; a Term Frequency–Inverse Document Frequency (TF–IDF) representation was additionally computed as implemented in the muon ATAC module, using the shifted log–transformation of term frequency (TF), which has been shown to be beneficial in sparse datasets compared with using TF directly as implemented in the muon ATAC module^5^.

To select highly variable genes, the scanpy function pp.compute_highly_variable was applied with flavor = ‘cellranger’ and n_top_genes = 3000. To select highly variable peaks, the scanpy function pp.compute_highly_variable was applied to the log–CPM–normalized ATAC data with parameters min_mean = 0.05 and min_disp = 0.5, resulting in 12,608 highly variable peaks. Cell cycle genes were scored to determine the cell cycle phase with the scanpy tl.score_genes_cell_cycle function per sample using the human S– and G2M–phase gene lists curated by Tirosh et al^6^.

#### Dimensionality reduction

For dimensionality reduction, PCA with 50 PCs was computed for gene expression, and for chromatin accessibility, we computed latent semantic indexing (LSI^7^), dropping the first component and retaining the next 30 dimensions, PCA on the highly variable peaks, and linear discriminant analysis (LDA) on the binarized peak matrix (see cisTopic^8^) as potential low–dimensional representations. To compute a joint UMAP projection of the multiome data, PCA representations were derived from highly variable features per modality, and the per-modality PCA embeddings were L2–normalized and used to compute a Weighted Nearest Neighbor (WNN) graph with 30 neighbors per modality; this joint graph was then used as input to the UMAP algorithm.

#### Batch correction

Batch correction was not performed in the final dataset because every batch correction method tested removed differences between the different subtypes of PDOs and left cell cycle variation as the dominant source of variation in the dataset, which is a clear sign of overcorrection.

#### Clustering and subtype characterization

Leiden^9^ clustering (leidenalg package *v.* 0.10.2) was first performed on the entire atlas to annotate broad categories. To identify subpopulations within each PDO, Leiden clustering was performed for each sample separately. For each sample, unimodal GEX and ATAC clustering were computed at resolution 1.0, and joint WNN clustering was computed at a resolution of 0.8. For the GEX clustering, gene characterization was performed with a one–vs–all t–test using the scanpy tl.rank_genes_groups function with method = ’t-test_overestim_var’. For differential accessibility analysis, considerable sparsity of the data was encountered; therefore, peak counts were aggregated using a pseudobulk approach. For each cluster and sample, we randomly sampled 80% of the cells and partitioned them into pseudobulk groups of 100 cells for the broad annotation or 20 cells for the subpopulation annotation. Pseudobulk counts were normalized with shifted log(CPM+1). We then characterized differentially accessible peaks using a one–vs–all Wilcoxon rank–sum test as implemented in the Muon package function atac.tl.rank_peaks_group (method = ’wilcoxon’, add_peak_type = True, add_distance = True, rank_abs = True). We then inspected the top 20 genes/peaks to assess whether a cluster represented a subpopulation. We repeated clustering and gene/peak characterization on the joint WNN clusters to label clusters. To set the subpopulation annotation in the context of the entire atlas, we performed another round of gene/peak characterization on each subpopulation with the entire PDO atlas as reference. Subpopulations were retained only when at least one of the top 20 characteristic genes/peaks was unique to the subpopulation; otherwise, they were merged with the majority label of the sample. The final annotation reflects a balance between the global context and the detailed resolution of subpopulations *per* sample.

#### Heterogeneity assessment

To assess group–wise heterogeneity, we derived a feature–centric metric as follows. Gene expression and chromatin accessibility were binarized per cell (X > 0). For each group (e.g., cluster or subtype), we counted the number of cells expressing a feature or accessible at a feature, normalized these counts across all features to obtain a relative abundance p_i, and computed the inverse Simpson Index (ISI)^10^ to quantify the diversity (D) of expressed genes and open peaks *per* group, respectively:

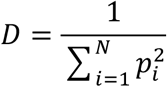

The ISI represents the number of effective features per group, such that a higher number represents higher diversity and therefore higher heterogeneity. To control for differences in sample size, we randomly sampled 1,000 cells per group and repeated draws 100 times; we report the mean ISI across draws. The ISI was computed independently over genes (GEX) and peaks (ATAC) at three groupings: main subtype (AR–high, AR–low, NEPC, WNT), refined subtype, and *per* PDO.

#### Subgroup analysis of gene expression modality

To perform a feature–based subgroup analysis of the gene expression modality, we used Spectra^11^ and Hotspot^12^. Briefly, we computed a Spectra model on the entire PDO atlas using the cross of PDO identity and the joint subtype annotation as the cell type annotation and using a subset of highly variable genes (parameters lam = 0.1, delta = 0.001, kappa = None, rho = 0.001, n_top_vals = 50, overlap_threshold = 0.2, gs_num = 3, num_epochs = 10,000). We used the ‘global’ gene–set dictionary as our *a priori* annotation dictionary and dropped all immune–related annotations. This resulted in 185 fitted factors. We then scored the gene expression for all factors using the scanpy tl.score_genes function and derived the mean *per* cell population, retaining factors whose maximum mean score across cell populations was ≥ 2; this yielded 32 factors representing 1,132 genes. To take into account the local correlation of genes, the Hotspot *v.* 1.1.1 compute_local_correlations function was applied to the retained gene set and grouped using the create_modules function in Hotspot (parameters min_gene_threshold = 30, core_only = True, fdr_threshold = 0.05), which split the genes into 15 gene modules. We then manually aggregated the 15 Hotspot modules into 6 modules by similarity in the local correlations plot. Per–module scores were computed with the scanpy tl.score_genes function, and cells were assigned to the module with the highest score. To interpret the modules, we ran over–representation analysis with the decoupler^13^ Python package *v.* 1.6.0 (Python v. 3.8.17) against the MSigDB Hallmark gene sets, with the background set to all genes retained after filtering genes present in fewer than 50 cells. Significantly enriched gene sets were filtered at FDR-adjusted p < 0.05.

#### Subgroup analysis of chromatin accessibility modality

To perform a feature–based subgroup analysis of the chromatin accessibility modality, we adapted the Latent Dirichlet Allocation (LDA^14^) workflow of the pycistopic package *v.* 1.10.12, Python 3.8.17. Due to performance limitations of the original pycistopic implementation, we transformed the binarized peak matrix into a bag–of–words representation, with each cell encoded as the list of accessible peak IDs, and replaced the python lda backend with the gensim^15^ LdaMulticore implementation. LDA is a probabilistic biclustering algorithm that groups peaks by co–accessibility to a predefined number of topics and simultaneously computes, for each cell, the probability of each topic, such that cells can be mapped to topics and peaks to topics. To select the number of topics *k*, we computed LDA models with k ranging from 2 to 24 topics and assessed each model by looking at the marginal probability distribution for all topics and topic overlap through normalized mutual information, as implemented in the normalized_mutual_info_score function of the scikit-learn^16^ package *v.* 1.3.0. We determined that k = 6 was the best number of topics to represent the data.

#### Motif enrichment analysis

For each annotated cluster, we performed differential accessibility testing in pseudobulks as described above. Peaks were retained when log-fold-change ≥ 0.6 and Benjamini-Hochberg-corrected p-value ≤ 10⁻⁵.

For each topic, we computed the standardized topic–region probability distribution for all peaks *per* topic and used the triangle thresholding method to determine the threshold to select peaks. Briefly, the triangle thresholding method is used on histograms where the histogram peak is connected with the furthest value at the histogram tail by a line, and the distance of the histogram values to this line is maximized. The histogram value where the maximum distance is observed represents the threshold.

We then used the run_pycistarget wrapper function from the SCENIC+3 package *v.* 1.0.1.dev3+g3741a4b for motif enrichment using both the Differentially Enriched Motifs (DEM) and the cisTarget (CTX) approaches, as implemented in pycistarget v. 1.0.3.dev1+g3fde1ce. We used the per–topic peak sets, with and without promoter regions, and the per–cluster differentially accessible peak sets at both the main–subtype and subclone resolution as the foreground set, and the set of all accessible peaks as the background set. This approach computes Cis–Regulatory Module (CRM) scores for foreground and background region sets, and the distribution of scores was compared using a Wilcoxon test. Motifs with a log–fold-change > 0.5 and a Bonferroni-corrected p-value < 0.05 were considered significant.

## RE–ANALYSIS OF HUMAN TUMOR SINGLE–CELL/SU2C AND NTP

### Re–analysis of human prostate single–cell RNA–seq data

We obtained pre–processed count matrices for all previously published patient samples in Zaidi et al.^17^, which have been deposited on GEO under the accession ID GSE264573 and https://duos.broadinstitute.org/ (accession no. DUOS-000115)^18^.

#### Quality Control (QC)

Similar to the QC of the single–cell multiomics data, we computed the number of total counts per cell, number of genes expressed per cell, and fraction of mitochondrial counts per cell. We defined the following permissive global thresholds to filter cells: library size between 500 and 100,000 UMIs, number of expressed genes between 200 and 12,000, and fraction of mitochondrial reads below 20%. In the second step, we computed the 2nd and 98th percentiles per sample for total counts per cell, number of genes expressed per cell, and fraction of mitochondrial counts to account for sample–to–sample variability in data quality. An exception was the fraction of mitochondrial reads, for which we did not use the 2nd percentile, as low fractions of mitochondrial reads do not present a disadvantage. Cells that passed all or all but one of the QC criteria were retained. After initial filtering, we detected doublets using the scrublet^2^ approach (v. 0.2.3) for gene expression data with parameters min_counts = 2, min_cells = 3, vscore_percentile = 85, n_pc = 50, expected_doublet_rate = 0.02, sim_doublet_ratio = 3, and n_neighbors = 15. The dataset after initial filtering consisted of 137,087 cells, which encompassed a mixture of tumor, stromal, and immune cells.

#### Normalization, feature selection and dimensionality reduction

We normalized the count matrix with shifted log–transformation of size-scaled counts. To select highly variable genes, we used the scanpy function pp.compute_highly_variable with parameters flavor = ‘cellranger’ and n_top_genes = 3000. We scored cell cycle genes to determine the cell cycle phase with the scanpy tl.score_genes_cell_cycle function per sample. As reference for cell cycle scoring, we used the human cell cycle gene list curated by Tirosh et al. For dimensionality reduction, we computed PCA with 20 components on the highly variable gene set using the scanpy pp.pca function and used this low-dimensional representation of the data to compute a k–nearest neighbor graph with k = 15 neighbors using the scanpy pp.neighbors function and a UMAP representation of the data using the scanpy tl.umap function.

### Clustering and cell type annotation

We performed Leiden^9^ clustering (leidenalg package *v.* 0.10.2) at resolution 0.5 with the scanpy tl.leiden function and computed the top 30 characteristic genes using the scanpy tl.rank_genes_groups function. We then examined these top genes per cluster and assigned clusters to broad compartments using marker genes for stroma (*COL1A2, ACTA2, PECAM1*), epithelial cells (*EPCAM*), neuroendocrine tumors (*SYP, CHGA, CHGB, ASCL1, NEUROD1*), and immune cells (pan–immune marker *PTPRC*). Within immune cells, we additionally distinguished T cells (*CD3D, IL7R, CD4, CD8A, CD69, CD38, LEF1, CCR7, TCF7, FOXP3*), NK cells (*GNLY, NKG7, KLRD1*), dendritic cells (*FCER1A, CST3*), B cells (*MS4A1, CD79A*), plasmablasts (*MZB1, HSP90B1, FNDC3B, PRDM1, IGKC, JCHAIN*), monocytes (*CD14, LYZ, FCGR3A, MS4A7*), and erythroid cells (*HBB, HBA2, HBA1*). Clusters without expression of known markers and with inconclusive characteristic gene expression were labeled as “unknown.”

#### Identification of tumor cells

To distinguish tumor cells from other epithelial and stromal cells, we predicted copy number variation (CNVs) using the Python implementation of inferCNV in the infercnvpy package *v.* 0.4.5 (https://infercnvpy.readthedocs.io/en/latest/) with reference categories Monocyte, T cell, B cell, Plasmablast, and Erythroid cells. Classification into normal and tumor tissue resulted in 33,379 tumor cells, which we used for subsequent analysis.

To identify tumor subsets, we recomputed the PCA and UMAP embedding of the data, repeated Leiden clustering at resolution 0.5, and computed the top 30 characteristic genes. We further examined the gene expression of tumor, immune, and myeloid marker genes as previously described in Zaidi et al.^17^ (*AR, KLK3, FOLH1, ALB, ASCL1, NEUROD1, PTPRC, LYZ, IGLC3*).

In accordance with the previously described analysis, we identified an *ALB*–expressing population, which is known to be expressed in hepatocytes. We further excluded two clusters with residual *LYZ* expression, known to be expressed in monocytes, and an *IGLC3* (Immunoglobulin Lambda Constant 3)–expressing cluster, indicative of B cell lineage cells. Lastly, we identified a cluster with cells from several patients that showed comparatively low library size and number of expressed genes. The resulting tumor dataset consisted of 30,105 cells from 11 patients (patient IDs HP7, HP8, HP10, HP11, HP13, HP14, HP17, HP18, HP19, HP20, and HP21).

#### Tumor subset classification

We first scaled the dataset by z–score and then used the NTP algorithm (see below) to classify cells by gene signatures using the top 500 differentially expressed genes of each major cell type lineage identified in the PDO atlas, using n_perms = 1000, BH FDR pooled across all comparisons, and assignment thresholds p_thresh = 0.05 and fdr_thresh = 0.5.

#### Re–Analysis of SU2C PRAD Cohort

The SU2C PRAD cohort bulk RNAseq dataset^19^ was downloaded from cBioPortal (https://www.cbioportal.org/study/summary?id=prad_su2c_2019) as FPKM–normalized expression. Values were z–scored gene–wise and classified with NTP using templates built from the top 500 differentially expressed genes of each major cell–type lineage identified in the PDO atlas, as described above.

#### SU2C Bulk RNAseq data processing

Raw sequencing reads were demultiplexed with bcl2fastq. Reads were trimmed with cutadapt *v.* 2.621 with parameters -j 8 -e 0.1 -q 20 -O 1 and aligned to build GRCm38 of the human genome using STAR v2.7.6a22^20^ with extra parameters --twopassMode Basic --outSAMunmapped Within --outSAMattributes NH HI AS NM MD --outSAMstrandField intronMotif. Alignments were quantified using all features in the Gencode version v29 M34 GTF23 with featureCounts from subread *v.* 1.6.224 in paired-end mode with options -O –fraction. Genes were then filtered down to those with at least 20 total counts across all samples across experiments.

### Classification of samples using gene signatures with the Nearest Template Prediction (NTP) algorithm

The classification of human tumor samples or single cells into molecular subtypes was performed using the Nearest Template Prediction (NTP) algorithm^21, 22^. This method allows for the prediction of classes in individual samples using a predefined set of marker genes (templates) without the need for a larger training cohort. We derived the gene signatures from the major lineages of the PDO atlas and used them to classify the SU2C PRAD cohort data^19^ and the human prostate cancer tumor biopsy single-cell RNAseq data^17^.

The algorithm proceeds in three stages:

1. **Template Construction:** A signature template is defined for each subtype using the top N differentially expressed genes identified from prior subcluster analysis. For each subtype, a binary template vector is created where genes specific to the subtype are assigned a value of 1, and all others are assigned 0.
2. **Distance Calculation:** For each sample, the proximity to each subtype template is evaluated. The algorithm calculates the Cosine distance between the sample’s actual expression profile and the ideal template vectors.
3. **Statistical Significance:** To ensure the robustness of the assignment, a permutation test is performed. A null distribution is generated by randomly resampling gene sets to calculate the probability (p-value) of obtaining the observed proximity by chance. False Discovery Rate (FDR) correction (Benjamini-Hochberg) is applied to account for multiple testing.

Samples were assigned to the subtype with the highest proximity, provided they met the predefined significance thresholds (e.g., p < 0.1 and FDR < 0.5). Samples failing to meet these criteria were labeled as "unclassified”.

We re-implemented the algorithm for compatibility with anndata v. 0.10.7, scanpy v. 1.10.0rc2.dev53+gefcf8df4, pandas v. 2.2.2, numpy v. 1.26.4 and statsmodels v.0.14.1 using Python v. 3.11.9.

## AHR CHIP–SEQ DATA

### Chromatin Immunoprecipitation (ChIP) and sequencing

Prostate cancer cells (∼4 million per condition) were provided to the MSKCC Epigenetics Research Innovation Lab for processing. Cells were fixed with 1% formaldehyde for 10 minutes, after which the reaction was quenched by the addition of glycine to a final concentration of 0.125 M. Fixed cells were washed twice with PBS, and cell pellets were resuspended in SDS buffer (100 mM NaCl, 50 mM Tris-HCl pH 8.0, 5 mM EDTA, 0.5% SDS, 1× protease inhibitor cocktail from Roche).

The resulting nuclei were pelleted and resuspended in immunoprecipitation buffer at 1 mL per 0.5 million cells. The immunoprecipitation buffer consisted of SDS buffer and Triton dilution buffer (100 mM NaCl, 100 mM Tris-HCl pH 8.0, 5 mM EDTA, 5% Triton X-100) mixed in a 2:1 ratio, supplemented with 1× protease inhibitor cocktail (Millipore Sigma, #11836170001). Samples were processed on a Covaris LE220+ focused ultrasonicator to achieve an average chromatin fragment length of 200–300 bp using the following parameters: PIP = 420, duty factor = 30, cycles per burst = 200, time = 35 minutes. Chromatin concentrations were estimated using the Pierce™ BCA Protein Assay Kit (Thermo Fisher Scientific, #23227) according to the manufacturer’s instructions. Immunoprecipitation reactions were set up in 1 mL of immunoprecipitation buffer in Protein LoBind tubes (Eppendorf, #22431081) and pre-cleared with 50 µL of Protein A/G magnetic beads (Cytiva, #9981922) for 2 hours at 4°C. After pre–clearing, samples were transferred to new Protein LoBind tubes and incubated overnight at 4°C with antibodies including AHR (rabbit monoclonal, Cell Signaling Technology, #83200) and H3K27ac (Cell Signaling Technology, #8173). The AHR antibody (CST #83200) is a rabbit monoclonal antibody recognizing the aryl hydrocarbon receptor (AHR), a ligand–activated transcription factor involved in xenobiotic metabolism and transcriptional regulation. It is validated for applications including ChIP, Western blotting, and immunofluorescence, and is used here to detect endogenous AHR chromatin binding. Antibodies were used at a dilution of 1:50 (AHR) or 1:100 (H3K27ac).The following day, 50 µL of BSA–blocked Protein A/G magnetic beads were added to each reaction and incubated for 2 hours at 4°C. Beads were then washed twice with low-salt wash buffer (150 mM NaCl, 1% Triton X-100, 0.1% SDS, 2 mM EDTA, 20 mM Tris-HCl pH 8.0), twice with high-salt wash buffer (500 mM NaCl, 1% Triton X-100, 0.1% SDS, 2 mM EDTA, 20 mM Tris-HCl pH 8.0), twice with LiCl wash buffer (250 mM LiCl, 10 mM Tris-HCl pH 8.0, 1 mM EDTA, 1% sodium deoxycholate, 1% IGEPAL CA-630), and once with TE buffer (10 mM Tris-HCl pH 8.0, 1 mM EDTA). Samples were reverse crosslinked overnight in elution buffer (1% SDS, 0.1 M NaHCO₃) and purified using the ChIP DNA Clean & Concentrator kit (Zymo Research, #D5205) following the manufacturer’s instructions. Recovered DNA fragments were quantified using a Qubit Flex fluorometer (Thermo Fisher Scientific) and submitted to the MSKCC Integrated Genomics Operation core facility for library preparation and sequencing. Immunoprecipitated DNA was quantified using PicoGreen, and fragment size distribution was assessed using an Agilent BioAnalyzer. Illumina sequencing libraries were prepared using the KAPA EvoPrep Kit (Roche, #10212250702) according to the manufacturer’s instructions, using 0.2–5 ng input DNA and 14 cycles of PCR amplification. Barcoded libraries were sequenced on a NovaSeq X plus in a paired-end 100 bp (PE100) run using the NovaSeq Reagent Kit (200 cycles) (Illumina). An average of 40–100 million paired-end reads were generated per sample.

#### ChIP–Seq Analysis

ChIP sequencing reads were trimmed and filtered for quality and adapter content using version 0.6.10 of TrimGalore (https://www.bioinformatics.babraham.ac.uk/projects/trim_galore), with a quality setting of 15, and running version 5.1 of cutadapt (https://cutadapt.readthedocs.io/en/stable/) and version 0.12.1 of FastQC (https://www.bioinformatics.babraham.ac.uk/projects/fastqc/). Reads were aligned to human assembly hg38 modified by 10X Genomics CellRanger software (version 2020-A) with bowtie2 (http://bowtie-bio.sourceforge.net/bowtie2/index.shtml) with parameters --local --very-sensitive --no-mixed --no-discordant --minins 10 --maxins 700 --dovetail and were deduplicated using MarkDuplicates in version 3.4.0 of Picard Tools (https://broadinstitute.github.io/picard/). To ascertain regions of chromatin accessibility, MACS3 version 3.0.3 (https://github.com/taoliu/MACS) was used with a q–value setting of 0.05 and scored against matched input as the control. The BEDTools suite version 2.31.1 (http://bedtools.readthedocs.io) was used to create normalized read density profiles. A global peak atlas was created by first removing blacklisted regions then merging all overlapping peaks and counting reads with version 2.1.1 of featureCounts. PyDESeq2^23^ was used to normalize read density (median ratio method) and to calculate differential enrichment for all pairwise contrasts. Peak–gene associations were computed by HOMER <u>annotatePeaks.pl</u> function version 4.11 with the cellranger-arc refdata-cellranger-arc-GRCh38-2020-A-2.0.0 genes.gtf file. Composite and tornado plots were created using deepTools v3.5.6^24^ by running computeMatrix reference-point and plotHeatmap on normalized bigwigs with flanking regions defined by the surrounding 1 kb from the peak center.

## ORGANOID CULTURE AND CRISPR SCREENS

### PDOs and culture

As described in Tang et al^25^, metastatic prostate cancer biopsies were collected with patient informed consent being obtained prior to tissue acquisition (Memorial Sloan Kettering Cancer Center, IRB 90–040, 06–107, and 16–865). Seventeen out of the 22 were previously published^25^. Patient–derived organoids (PDOs) were established and cultured as previously described^25^. Organoids were passaged every 4–7 days based on growth rates by enzymatic dissociation using TrypLE (Gibco) at 37 °C until a single–cell suspension was obtained. Digestion was supplemented with Y-27632 (10 μM) to inhibit anoikis. All lines were mycoplasma free by PCR testing.

#### PDO H&E and Immunofluorescence

Haematoxylin and eosin (H&E) staining was performed as previously described. Automated multiplex immunofluorescence (IF) staining was conducted using the Leica Bond BX staining system. Paraffin–embedded tissues were sectioned at 5 μm and baked at 58 °C for 1 h. Slides were loaded onto the Leica Bond platform, where IF staining was performed as follows: samples were dewaxed at 72 °C and pretreated with EDTA–based epitope retrieval ER2 solution (Leica, AR9640) for 20 min at 100 °C. Triple or quadruple antibody staining and detection were performed sequentially. Primary antibodies were incubated for 1 h at room temperature, followed by incubation with Leica Bond Polymer anti–rabbit HRP secondary antibody (included in the Polymer Refine Detection Kit, Leica, DS9800) for 8 min at room temperature. For mouse primary antibodies, a rabbit anti–mouse linker (Leica Bond Post-Primary reagent, included in the Polymer Refine Detection Kit) was applied for 8 min before incubation with the anti-rabbit HRP polymer. Signal detection was performed using Alexa Fluor tyramide signal amplification reagents (Thermo Fisher Scientific, B40953, B40958) or CF® dye tyramide conjugates (Biotium, 92174). Complete antibody information is provided in **Supplementary Table 14.** After each round of staining, epitope retrieval was repeated to denature primary and secondary antibodies before application of the next primary antibody. Upon completion of staining, slides were washed in PBS and incubated with 4′,6–diamidino–2–phenylindole (DAPI; 5 μg/ml; Sigma-Aldrich) for 5 min, rinsed in PBS, and mounted using Mowiol 4–88 (Calbiochem). Slides were stored overnight at −20 °C before imaging.

#### PDO Clonality

Organoid culture and seeding experiments were performed as previously described^18, 25^. During seeding assays, organoid medium was supplemented with the anoikis inhibitor Y-27632 (10 μM). For outgrowth quantification, 100 cells were seeded per well in triplicate per condition, and organoids were counted 7 days post–seeding as three independent replicates.

#### PDO Cas9

Patient–derived organoids (PDOs) were transduced with Cas9 cDNA and maintained under blasticidin selection (5–15µg/mL).

#### Mini–pool creation RNA-seq and Differential Expression Analysis

RNA sequencing data from PDOs were analyzed in R (v4) using the DESeq2 package. Raw gene–level counts were imported, and samples were assigned to the prior bulk–derived conditions from Tang et al.: AR, NEPC, WNT, and SCL. Genes with total counts of 20 or fewer across all samples were excluded prior to analysis. Count data were normalized using variance–stabilizing transformation (VST). Differential expression between conditions was assessed using the Wald test. P values were adjusted for multiple testing using the Benjamini–Hochberg method. Genes with an adjusted P value < 0.05 and absolute log2 fold change > 1.5 were considered significantly differentially expressed. For more permissive threshold for TFs at lower expression, we applied a threshold of log2FC >= 0.43, FDR <0.05.

#### Methods for sgRNA library design and cloning

sgRNA sequences were identified using CRISPick (https://portals.broadinstitute.org/gppx/crispick/public), selecting the top four sgRNAs per gene based on Pick Order ranking criteria. sgRNA sequences were divided into three sub–pools, and sgRNA oligonucleotides were synthesized on–chip (Agilent) as previously described^26^. Synthesized oligos were PCR–amplified, and amplicons were cloned via Golden Gate assembly into BsmBI restriction sites of the pUSEPR backbone^27^. Cloned libraries were amplified by electroporation into electrocompetent cells (Lucigen), and plasmid DNA was isolated using Qiagen kits. Library representation was assessed by next–generation sequencing (Illumina). List of sgRNA and/or library QC HiSeq metrics are shown in **Supplementary Table 4 and Table 6**.

#### Lentiviral production

Lentiviruses were produced by co–transfection of HEK293T cells (ATCC) with lentiviral backbone constructs and packaging vectors (psPAX2 and pMD2.G; Addgene 12260 and 12259) using TransIT–LT1 (Mirus Bio, MR 2306). Virus was concentrated using Lenti–X concentrator from Takara Bio.

#### Infection of Minipools Library 1–3 and Comprehensive Libraries

All transductions were performed by spin–infection at 1,500 r.p.m. for 1.5–2 h at room temperature. For each patient-derived organoid (PDO), cells were expanded to maintain a coverage of 8,000×. Cells were spin–infected with each given library at a multiplicity of infection (MOI) of 0.3–0.5 in 6–well plates using viral supernatant. Viral titers for each cell line were determined by flow cytometry by calculating the percentage of TagRFP–positive cells relative to total cells. Following infection, cells were selected with puromycin for 3–4 days, after which half of the cells were harvested and frozen as the initial time point (T0), and the remaining cells were reseeded to maintain 8,000× representation. Cells were subsequently cultured and harvested at 21 and 28 days post–infection for DNA extraction.

#### DNA extraction and sequencing

Cell pellets were lysed, and genomic DNA was extracted using Qiagen kits and quantified by Qubit (Thermo Fisher Scientific). A quantity of genomic DNA corresponding to 5,000–8,000× representation of sgRNAs was PCR–amplified to add Illumina adapters and multiplexing barcodes. Amplicons were quantified by Qubit and Bioanalyzer (Agilent) and sequenced on an Illumina HiSeq 2500 platform. Sequencing reads were aligned to the screened library, and counts were obtained for each sgRNA.

#### Methods for MAGeCK analysis

FASTQ pre–processing and guide abundance were determined using the MAGeCK count command. The 5′ trim length was automatically detected by MAGeCK, and a normalized count file was generated using median normalization. Sample quality control showed greater than 85% mapped reads and a minimum sample correlation of 0.8 across all replicates. Enrichment was determined using the MAGeCK RRA (robust rank aggregation) function to obtain gene–level enrichment scores, and P values were determined by permutation. All analyses were performed using MAGeCK *v*0.5.9. All MAGeCK results are shown in **Supplementary Table 5 and Table 7**.

#### sgRNA depletion experiment and competition assay

Sequences for guide RNAs of interest were cloned into the pUSEPR backbone and verified by Sanger sequencing. A total of 2–4 × 10⁵ cells from organoid lines stably expressing Cas9 were transduced with sgRNAs at a multiplicity of infection (MOI) of 0.3–0.4. At indicated time points, 2 × 10⁵ cells were split and reseeded in 12-well plates, and the remaining cells were collected for flow cytometry analysis. Cells were imaged for TagRFP and DAPI signals at days 4, 11, 18, 25 and 32, or at days 4, 8, 16, 23, 30, 37 and 43. Data acquisition was performed using a FACSymphony A3 (BD Biosciences), and values were normalized to day 4 measurements.

#### PCA15 TROP2 and NCAM1 Flow Cytometry and FACS

PDO PCA15 was selected for lineage reconstitution experiments based on the presence of admixed adenocarcinoma/SCL, intermediate, and NEPC populations identified by single–cell multiome profiling. Organoids were dissociated into single cells using TrypLE Express (Gibco) supplemented with Y–27632 ROCK inhibitor (10 µM), stained for 30 minutes on ice in PBS containing 0.05% BSA and Y–27632 (10 µM) using TROP2–PE (BD Cat. 564837, Clone: 162-46, 1:100) and NCAM1–Alexa647 (BD Cat. 557711, Clone B159, 1:100). DAPI was used as a viability dye to exclude dead cells. Flow cytometric sorting was performed to isolate TROP2 single–positive (adenocarcinoma/SCL/ASCL1–low), TROP2/NCAM1 double–positive (intermediate), and NCAM1 single–positive (NEPC–A/N) populations, alongside an unsorted parental population control. Following sorting, cells were reseeded in Matrigel and cultured under standard PDO conditions for 3 weeks to assess lineage reconstitution potential. At endpoint, organoids were dissociated again into single cells using TrypLE Express supplemented with Y–27632 (10 µM), restained with TROP2–PE and NCAM1–Alexa647, and analyzed by flow cytometry to evaluate the ability of each sorted population to regenerate different lineages.

## INTEGRATION OF CRISPR AND SC–MULTIOME DATA

### Subpopulation Impact Analysis

We integrated CRISPR screen results (discussed below, beta values) with single–cell multiome data aggregated at two levels of resolution: the PDO level and the subclone level. The analysis was restricted to 5 PDOs that have both pooled CRISPR screen data and a non-trivial subclone structure. For the CRISPR outcomes, we used two different significance thresholds: p < 0.1 for mini-libraries and p < 0.2 for genome-scale libraries.

#### Global Correlation Analysis

To identify transcription factors (TFs) whose dependencies were consistently associated with molecular states across different models, we calculated the Pearson correlation coefficient between the binarized CRISPR beta values and the mean molecular features (Expression and Activity) for each gene across all PDOs. TFs were classified as “high–correlation” if the absolute correlation coefficient exceeded 0.5.

### Leave-One-Out (L1O) Linear Regression Framework

To systematically assess how well bulk molecular features predict CRISPR outcomes compared to subpopulation features, we implemented a Leave-One-Out (L1O) linear regression procedure:

`o` **Baseline Fit:** For each gene, a linear regression model was trained using the beta values as the independent variable to predict the molecular feature (𝑦_observed_ = mean gene expression or activity) across all PDOs.
`o` **L1O Procedure:** For each specific PDO that has a subclone structure, we refit the linear model using data from all other PDOs. This “chimera” model was then used to predict the molecular feature 𝑦of the held–out PDO.
`o` **Residual Computation:** We calculated the residuals as the difference between the observed molecular feature and the predicted value (𝑦) derived from the L1O model:

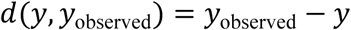

- **Relative change in** 𝑅^2^: For each gene and PDO, we computed the relative change in the coefficient of determination (𝑅^2^) when the held-out PDO or one of its subpopulations was virtually included in the regression without refitting the model:

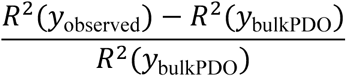

**Evaluating Subpopulation Improvement.** We evaluated whether molecular data from specific subclones provided a better fit for the observed CRISPR depletion than the bulk PDO data.

- **Proportional Improvement in Residual:** We calculated the relative reduction in the absolute residual when replacing the bulk PDO value with a subpopulation–specific value:

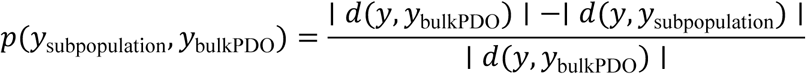

A positive value indicates that the subpopulation molecular state more closely aligns with the linear trend established by the rest of the dataset/

- 𝑅^2^**Optimization (“Winning Subpopulation” Analysis):** For each gene and PDO, we identified the “winning subpopulation” — the subpopulation population that maximized the coefficient of determination (𝑅^2^) when included in the regression. We then refit the global linear model using these optimal subpopulation features to determine the “Best 𝑅^2^,” comparing it against the “Baseline 𝑅^2^” (derived from the baseline model described above) to quantify the total gain in explanatory power provided by accounting for subpopulation heterogeneity.

#### Statistical Independence Testing

To determine whether the improvement in model fit was driven by molecular features of the specific subpopulations, such as a subpopulation having the lowest expression, we employed a G–test, a likelihood–ratio test. This test evaluated the independence between the subpopulation with the minimum feature value and the su subpopulation bclone providing the maximum 𝑅^2^improvement across all genes and PDOs.

## AHR STUDIES

### KYN-101 studies

A total of 0.04 × 10⁶ cells per well were plated with or without 1 μM KYN-101(MedChemExpress, Cat #HY–134217) in triplicate in a 24–well plate format. Cells were counted, split, and re–treated with drug every 7 days for a total duration of 21 days. Cell counts were adjusted to account for reseeding fractions used for continued culture and scaled linearly to reflect the expected cell numbers if the entire culture had been maintained.

#### qPCR

RNA was extracted from cells using the RNeasy Mini Kit (Qiagen) according to the manufacturer’s instructions. Isolated RNA was quantified using a NanoDrop spectrophotometer and stored at −80 °C. cDNA was synthesized from 500 ng of RNA using the LunaScript RT SuperMix Kit (New England Biolabs) according to the manufacturer’s protocol and stored at −20 °C. Quantitative PCR (qPCR) was performed using 1 μl of cDNA diluted 1:10, 5 μl of SsoAdvanced Universal SYBR Green Supermix (Bio-Rad), and 0.5 μl of PrimePCR primers (Bio-Rad) in a total reaction volume of 10 μl. Primers were designed to span introns for target genes. Reactions were run in technical triplicate on a ViiA 7 Real-Time PCR System (Thermo Fisher Scientific). Gene expression changes were calculated using the 2^−ΔΔCt^ method.

#### Western blot

PDOs were treated with cell recovery solution (Corning), incubated, and washed with PBS to remove Matrigel. Cells were lysed in 1× RIPA lysis buffer (Millipore) supplemented with protease and phosphatase inhibitors, with vortexing every 3 min for a total of 30 min. Lysates were clarified by centrifugation at maximum speed for 10 min at 4 °C. Protein concentration was determined using the Pierce BCA Protein Assay Kit (Thermo Fisher Scientific). Equal amounts of protein were mixed with 1× Laemmli sample buffer (Bio-Rad) and boiled for 10 min before separation by SDS–PAGE on 4–15% Mini-PROTEAN TGX Stain-Free gels (Bio-Rad). Proteins were transferred onto PVDF membranes (Bio-Rad), and a Full Range Rainbow recombinant protein ladder (Cytiva) was used as a molecular weight marker. Membranes were blocked in 5% non-fat dry milk in 0.1% Tween-20 TBST for 1 h at room temperature with agitation and incubated overnight at 4 °C with primary antibodies diluted 1:1,000 in 5% milk. Membranes were washed three times for 15 min in 0.1% Tween-20 TBST and then incubated with secondary antibodies diluted 1:5,000 in 5% milk for 1 h at room temperature. Membranes were subsequently washed repeatedly over 30 min in 0.1% Tween-20 TBST. Signal was detected using ECL Prime Western Blotting Detection Reagent (Cytiva) and imaged on a ChemiDoc imaging system (Bio-Rad).

## Data and code availability

Next–generation sequencing data have been deposited at GEO and will be publicly available as of the date of peer–reviewed publication.

