## Supplementary Figures and Legends for "Dependencies in heterogeneous, lineage plastic patient–derived prostate cancer organoids revealed through integrated single–cell multiomics and CRISPR screening"

Supplementary Figure 1

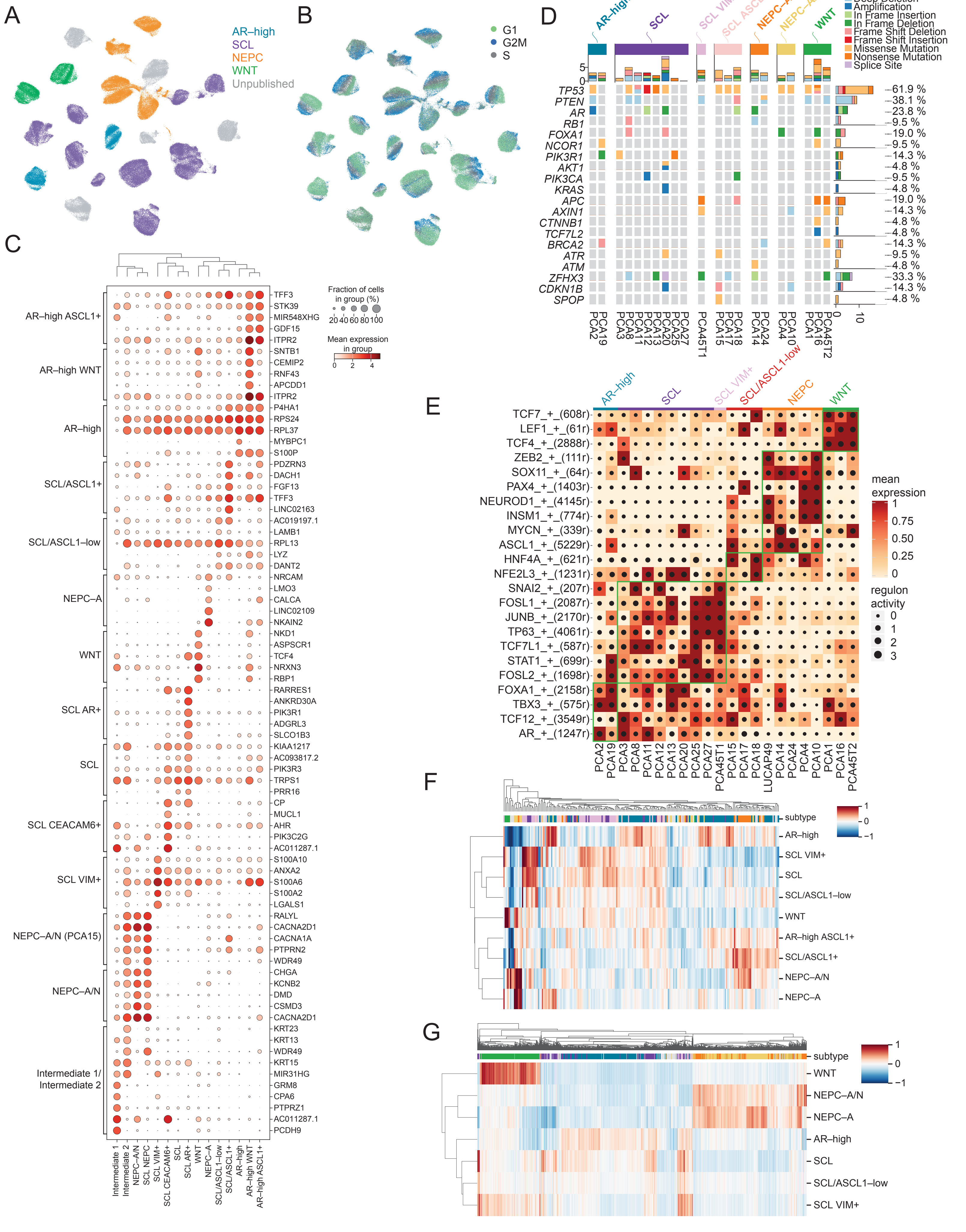

### Supplementary Figure 2

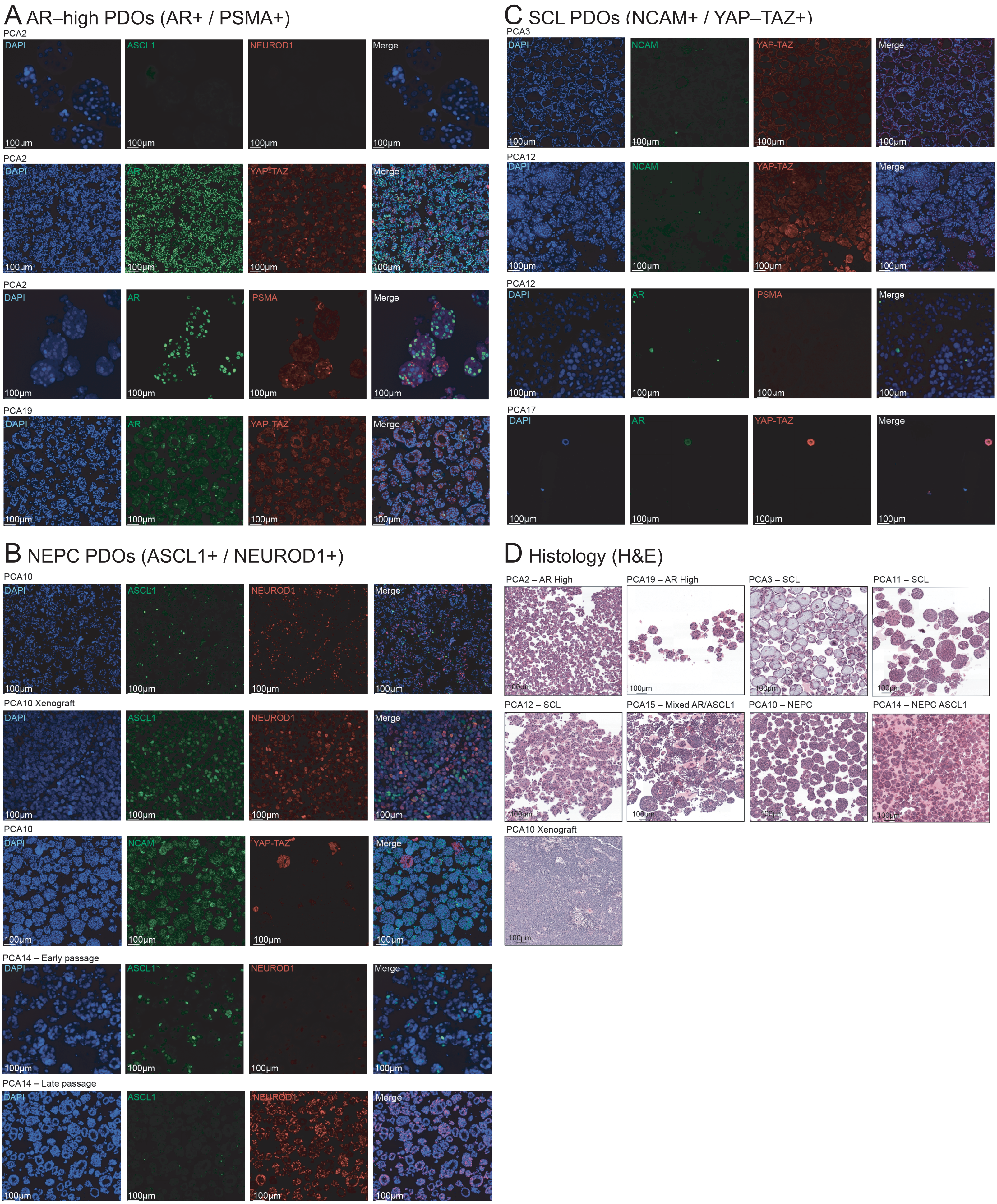

Supplementary Figure 3

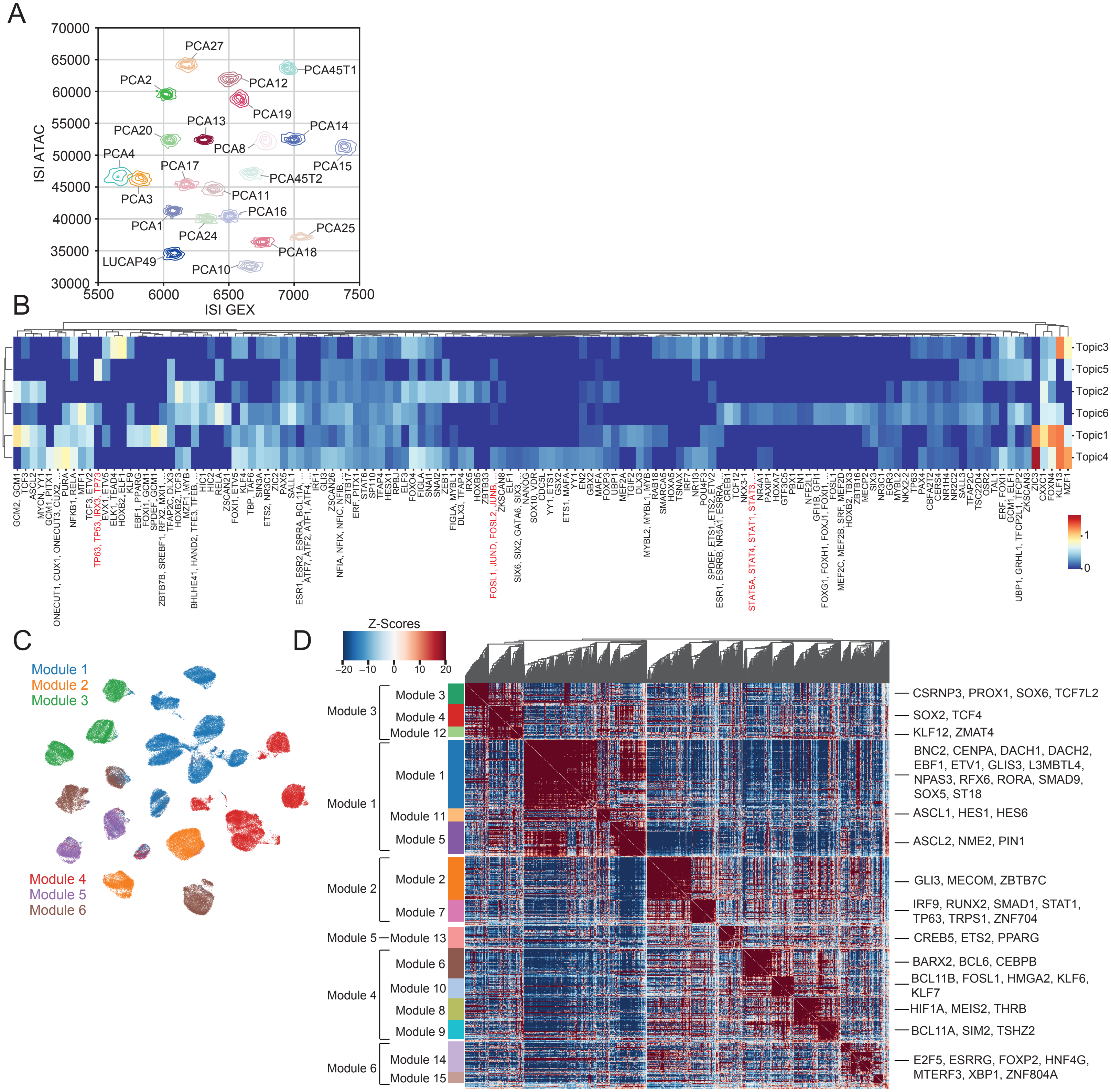

### Supplementary Figure 4

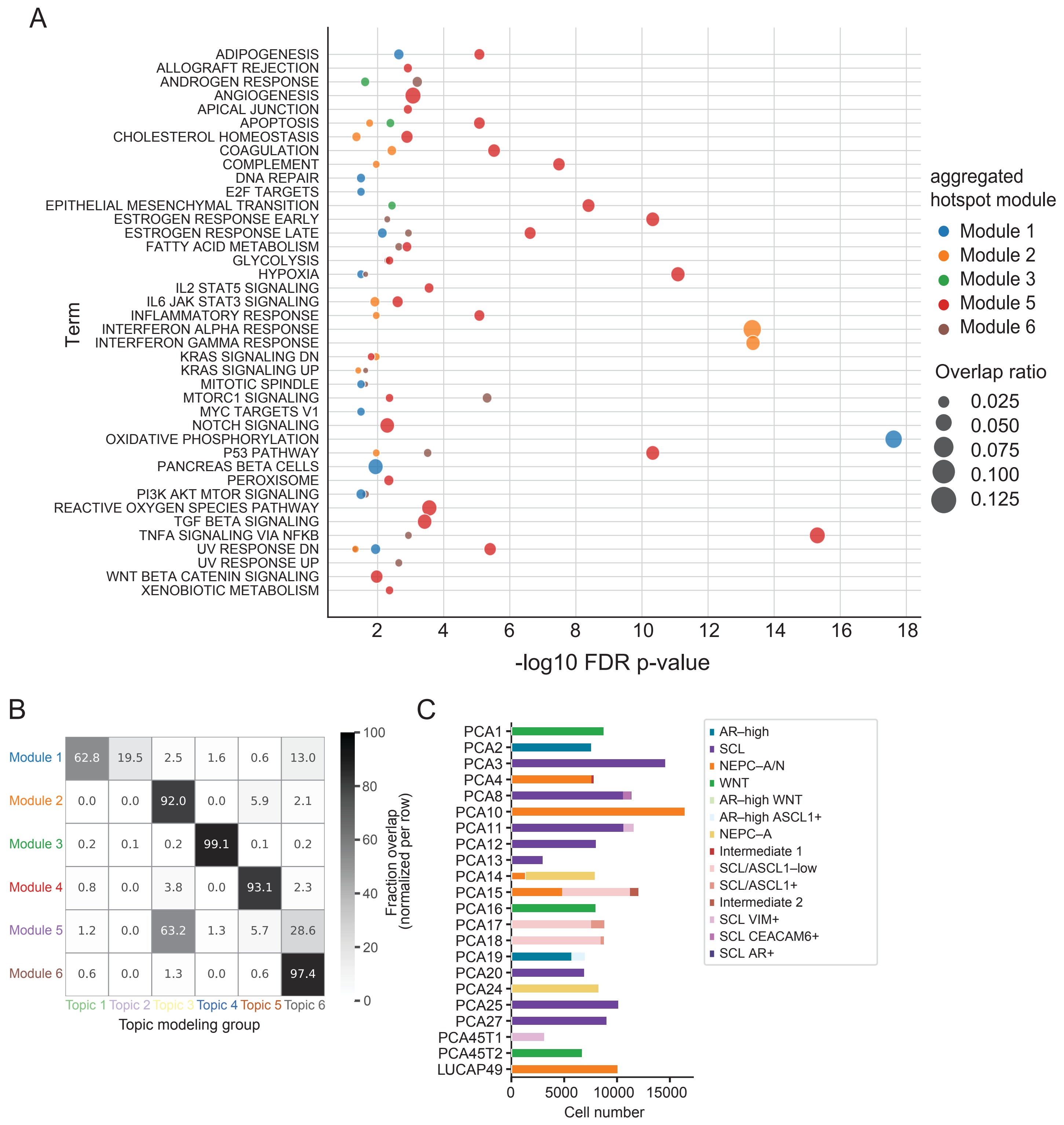

### Supplementary Figure 5

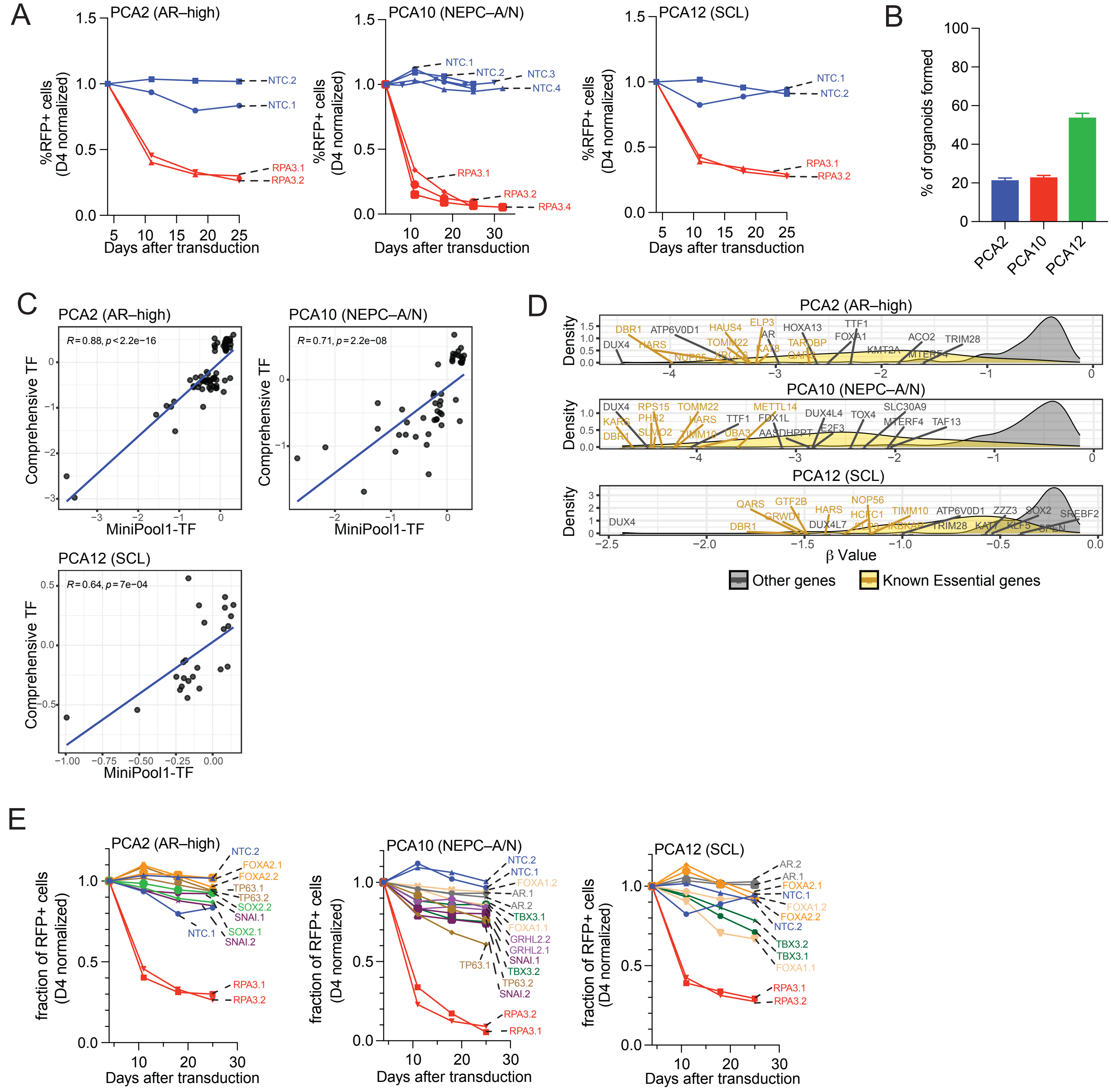

### Supplementary Figure 6

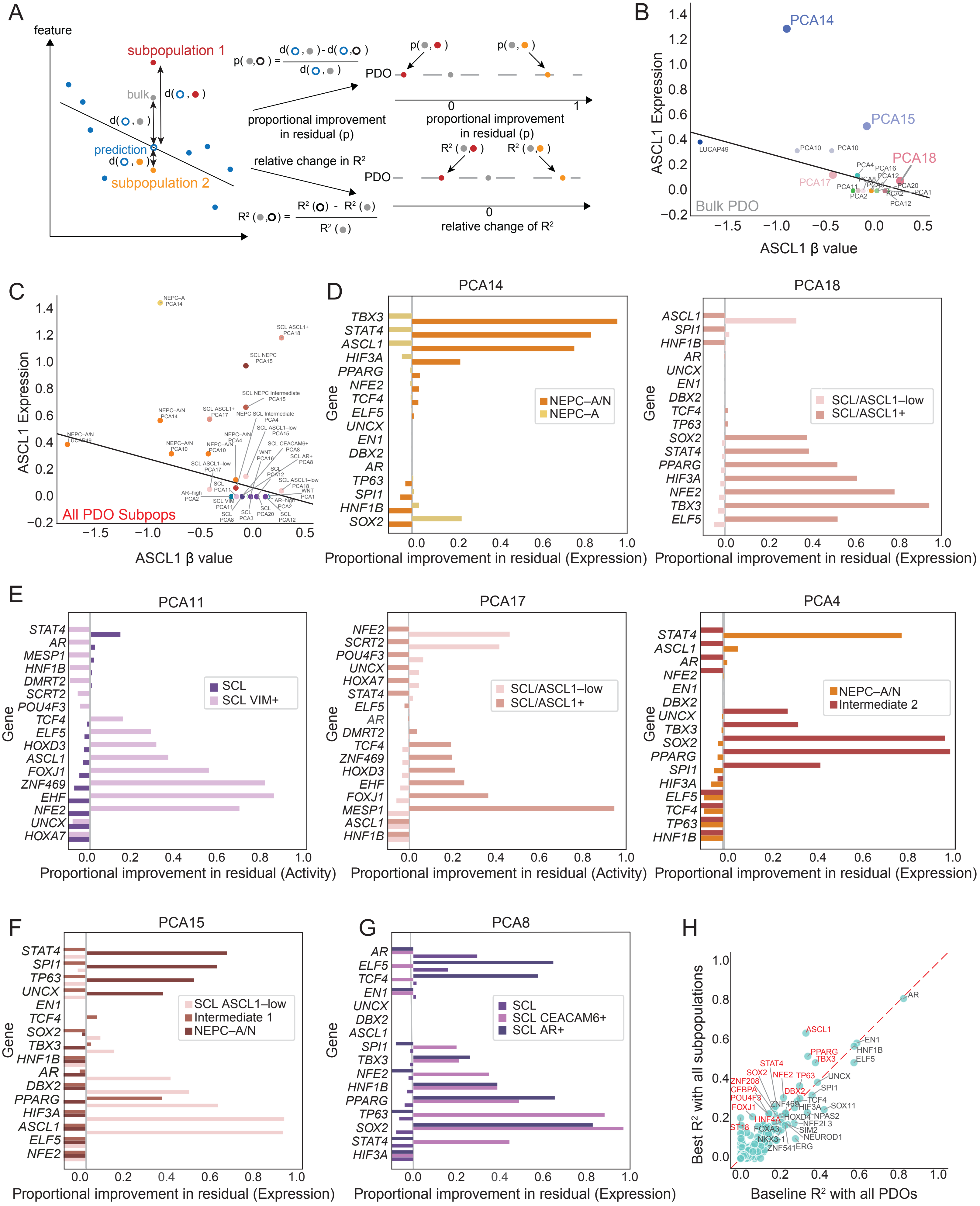

### Supplementary Figure 7

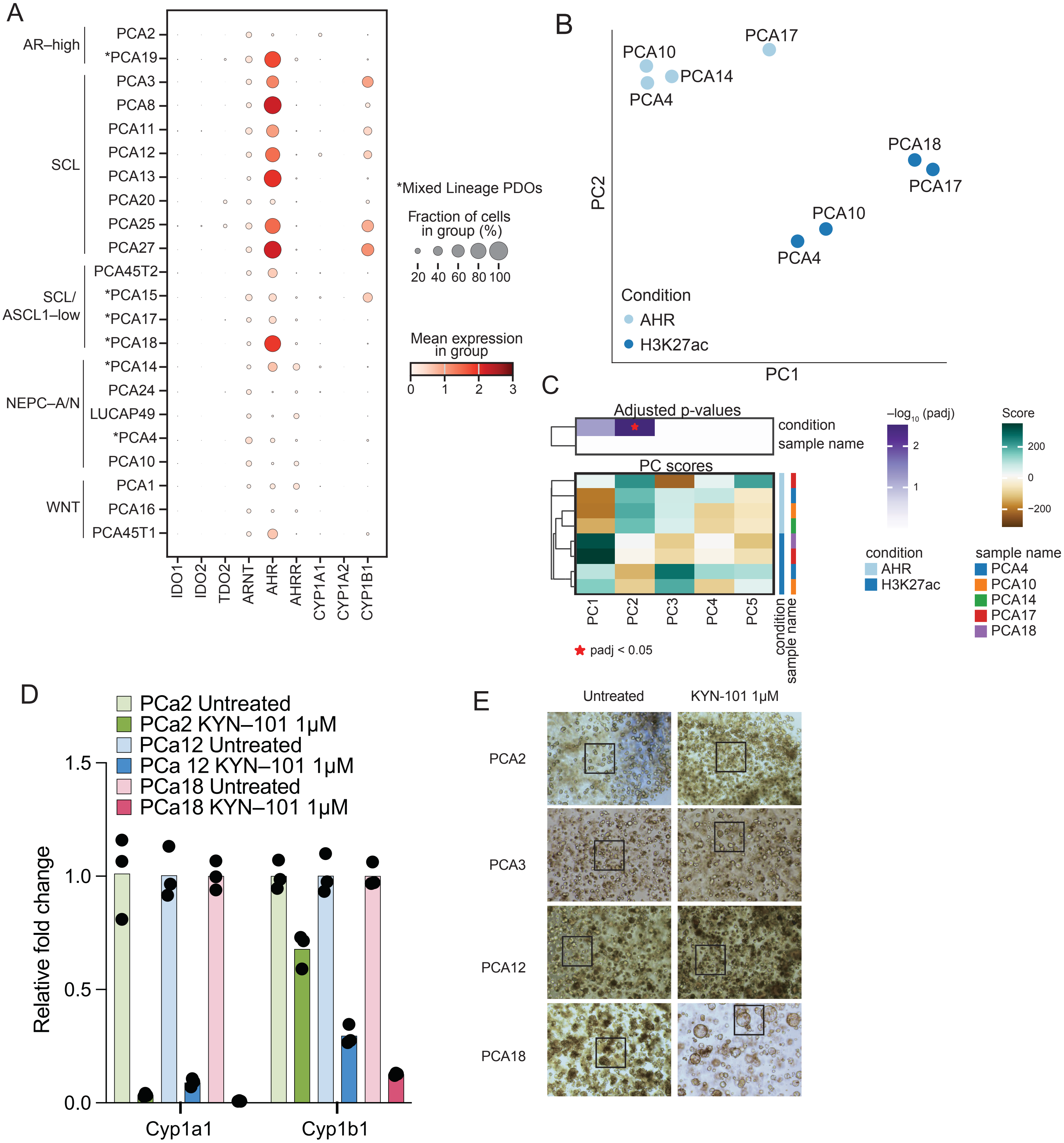

#### Supplementary Figure Legends

**Supplementary Figure 1. Single–cell multiome–based subtype definition and genomic features of patient–derived organoids (PDOs). (A)** UMAP projection of single–cell RNA–seq and ATAC–seq data from 22 PDOs colored by previously bulk–defined subtypes [PMID: 35617398]. PDOs labeled in grey are unpublished. **(B)** UMAP of scored cell cycle phase *per* cell across PDOs. **(C)** Dot plot showing the fraction of cells and mean expression of representative marker genes across refined lineage states and intermediate populations. Dot sizes indicate fraction of cells in each group and colors indicate mean expression. **(D)** Oncoprint showing genomic alterations in recurrent prostate cancer genes across PDOs, grouped by subtype. Alteration classes are indicated by color. **(E)** Heatmap of SCENIC+ regulon activity per PDO, highlighting subtype–associated transcription factor programs. **(F)** Application of PDO–derived subtype signatures to bulk transcriptomic data from the Stand Up To Cancer (SU2C) metastatic castration–resistant prostate cancer cohort. **(G)** Application of PDO–derived subtype signatures to independent single–cell RNA–seq data from human prostate cancer [PMID: 38968122]

**Supplementary Figure 2. Immunofluorescence across patient-derived organoids (PDOs) lineages. (A)** Immunofluorescence staining of AR–high PDOs showing strong expression of androgen receptor (AR) in PCA2, but focally in a subset of cells in PCA19. **(B)** Immunofluorescence staining of neuroendocrine prostate cancer (NEPC) PDOs showing expression of neuroendocrine markers ASCL1 and NEUROD1. **(C)** Immunofluorescence staining of SCL PDOs showing NCAM and activated YAP/TAZ expression. **(D)** Representative hematoxylin and eosin (H&E) staining of PDO–derived organoids and PCA10 Xenograft.

**Supplementary Figure 3. Quantification of regulatory heterogeneity and identification of chromatin accessibility programs. (A)** Inverse Simpson Index (ISI) for gene expression (GEX) and chromatin accessibility (ATAC) *per* PDO. **(B)** Identification of chromatin accessibility "Topic Modules" using latent Dirichlet allocation (LDA), with associated transcription factor enrichments. **(C)** UMAP projection of single–cell data colored by Spectra–derived "Hotspot Modules" identified from highly variable genes. **(D)** Correlation heatmap of genes assigned to Hotspot Modules derived from Spectra matrix factorization. Modules are hierarchically clustered based on gene–gene correlations, and representative transcription factors associated with each module are indicated. Color scale represents Z–scores.

**Supplementary Figure 4. Integration of chromatin accessibility and gene expression programs. (A)** Bubble plot showing enrichment of Hallmark gene sets across aggregated Hotspot Modules identified by Spectra matrix factorization. The x-axis indicates –log10 False Discovery Rate (FDR)–adjusted p–values, and dot size represents overlap ratio. Colors denote module identity. **(B)** Overlap between Topic Modules and Hotspot gene expression modules, showing concordance between chromatin accessibility topic and transcriptional module programs. **(C)** Stacked bar plot showing the number of cells contributed by each PDO, colored by refined subtype identity.

**Supplementary Figure 5. Validation and benchmarking of CRISPR screening in patient-derived organoids (PDOs). (A)** Fraction of red fluorescent protein (RFP)–positive cells over time following transduction of PCA2 (AR-high), PCA10 (NEPC-A/N), and PCA12 (SCL) PDOs with non-targeting control (NTC) sgRNAs or sgRNAs targeting the essential gene RPA3. Values are normalized to day 4 after transduction. **(B)** Clonality assay showing the percentage of organoids formed in PCA2, PCA10 and PCA12 PDOs one week after seeding 100 single cells. **(C)** Correlation of gene-level depletion (β values) between mini-pool transcription factor (Mini Pool 1–TF) and comprehensive CRISPR screens in PCA2 (AR–high), PCA10 (NEPC–A/N), and PCA12 (SCL) PDOs. Each point represents a gene. Pearson correlation coefficients (*R*) and corresponding P–values are shown. **(D)** Distribution of gene–level depletion scores (β values) for known essential genes (yellow) compared to all other genes (gray) across PCA2, PCA10, and PCA12 PDOs. Density plots show stronger depletion of essential genes (yellow). Selected genes are annotated. **(E)** Fraction of RFP–positive cells over time following transduction of PCA2 (AR–high), PCA10 (NEPC–A/N), and PCA12 (SCL) PDOs with sgRNAs targeting selected genes from CRISPR screen, alongside non–targeting control (NTC) and the essential gene control RPA3. Values are normalized to day 4 after transduction. Multiple sgRNAs per gene are shown.

**Supplementary Figure 6. Subpopulation–resolved modeling of gene dependency in heterogeneous PDOs. (A)** Schematic illustrating the framework used to evaluate gene dependency modeling at bulk *versus* subpopulation levels. **(B)** Relationship between *ASCL1* gene dependency (β value) and gene expression across PDOs. Each point represents a PDO, with selected samples annotated. **(C)** Relationship between *ASCL1* gene dependency (β value) and gene expression across all PDO subpopulations. Each point represents a subpopulation, highlighting increased heterogeneity compared to bulk–level analysis. **(D)** Proportional improvement in residuals for prediction of gene expression across subpopulations compared to bulk for selected transcription factors in representative PDOs (PCA14 and PCA18). Bars are stratified by subpopulation identity. **(E–G)** Proportional improvement in residuals for prediction of transcription factor activity or expression across subpopulations compared to bulk in representative PDOs (PCA11, PCA17, PCA4, PCA15, and PCA8). Bars are grouped by subpopulation identity. **(H)** Comparison of baseline $R^2$ (bulk model) versus best $R^2$ obtained using subpopulation–resolved modeling across transcription factors. Each point represents a gene; selected transcription factors are annotated. The dashed line indicates equality.

**Supplementary Figure 7. Aryl hydrocarbon receptor (AHR) pathway activity and pharmacologic inhibition across prostate cancer patient–derived organoids (PDO) subtypes.** **(A)** Dot plot showing expression of AHR pathway genes across PDOs stratified by lineage subtype. Dot size represents the fraction of cells expressing each gene; colors indicate mean expression level. AHR pathway activity is enriched in specific SCL, SCL VIM+, SCL/ASCL1–low PDOs. **(B)** Principal component analysis (PCA) of chromatin accessibility profiles comparing untreated and AHR-inhibited (KYN-101, 1µM) conditions across PDOs (PCA4, PCA10, PCA14, PCA17, PCA18). **(C)** Heatmap of principal component (PC) scores and corresponding adjusted p values across PDOs and treatment conditions (AHR inhibitor versus control). Significant changes ($p_{adj} < 0.05$; red asterisk) highlight treatment–associated shifts in chromatin state. **(D)** Quantitative polymerase chain reaction (qPCR) analysis of AHR target genes in representative PDOs following treatment with KYN–101 (1 µM) compared to untreated controls at 7 days after treatment. Data are shown as relative fold change normalized to control conditions; points represent biological replicates. **(E)** Representative brightfield images of PDOs treated with vehicle or KYN-101 (1 µM), illustrating morphological changes and growth effects in PCA18.
